# The Fork Protection Complex protects long replicons from DNA damage at the cost of genome instability induced by DNA topological stress

**DOI:** 10.1101/2023.08.04.551986

**Authors:** Andrea Keszthelyi, Alex Whale, Jon Houseley, Jonathan Baxter

**Affiliations:** Genome Damage and Stability Centre, School of Life Sciences, Science Park Road, University of Sussex, Falmer, Brighton, East Sussex BN1 9RQ, U.K.; The Babraham Institute, Babraham Research Campus Cambridge, CB22 3AT

## Abstract

Tof1/Timeless protects eukaryotic cells from DNA replication stress as part of the Fork Protection Complex (FPC). Tof1 supports rapid DNA replication, fork pausing, and resolution of DNA topological stress. Here, we show that disruption of FPC function through loss of either Tof1 or Mrc1 results in DNA damage in long replicons. Despite increasing DNA damage in long replicons, loss of either Tof1 or Mrc1 concurrently reduces DNA damage in regions prone to damage caused by DNA topological stress, indicating that the rapid replication promoted by the FPC fosters completing DNA replication at the cost of increased vulnerability to DNA topological stress. Supporting this we find that a *tof1* mutation that selectively inhibits DNA topological stress resolution increases DNA damage in contexts prone to DNA topological stress. Our data indicates that the FPC balances rapid replication with recruitment of topoisomerase I to resolve the topological stress generated by increased DNA unwinding.

## INTRODUCTION

The faithful replication of DNA is hindered by numerous endogenous and exogenous factors, collectively termed replication stress ^1^. Certain chromosomal sites, termed fragile sites, are constitutively enriched for DNA damage markers due to replication stress caused by site specific DNA and protein structures ^2, 3^. Whilst some genomic contexts constitutively induce replication stress, other fragile sites are only revealed following perturbation of replication dynamics. For example, low doses of the polymerase inhibiting agents reveal a subset of fragile sites in chromosomes, often linked to unusually long replicon distance ^4^.

The evolutionarily conserved fork protection complex (FPC) has multiple functions in promoting faithful DNA replication, including enabling rapid elongation of unstressed forks, stabilising the replisome under conditions of replication stress and mediating DNA replication checkpoint signalling ^5, 6^. FPC function has primarily been investigated in the budding yeast *Saccharomyces cerevisiae* (*S.c.*), the fission yeast *Schizosaccharomyces pombe* (*S.p.*) and human (*H.s.*) cell lines. In *S.c.* the three FPC proteins are called Tof1, Csm3 and Mrc1, in *S.p.* Swi1, Swi3 and Mrc1 and in *H.s.* Timeless, Tipin and Claspin. The FPC is located at the front of the replisome and maintains multiple conserved interactions within the replisome, with reported contacts and interactions between FPC factors and MCMs 2, 4, 6, 7, Cdc45, AND-1, Pol Epsilon, Top1, DDX11, Rpa1, Cdc7, PARP1, SDE2, and Spt16 ^7–18^. The FPC also contacts the parental duplex DNA and alternate DNA (i.e., G4) conformations ^7, 19^.

The FPC can be considered as two partially independent units, Tof1-Csm3 and Mrc1. Mrc1 mediates replication fork speed, signalling of the replication checkpoint to inhibit cells from traversing mitosis, preventing deleterious fork processing and stimulating local DNA damage repair ^20–24^. *In vivo* and *in vitro* evidence indicates that Tof1-Csm3 stabilise Mrc1 interactions with the replisome, thus facilitating Mrc1 function. In contrast, Tof1-Csm3 associates with the replisome independently of Mrc1 and supports some functions independently of Mrc1. Tof1-Csm3 (Swi1-Swi3 in *S. pombe*) promote fork pausing at a variety of protein-DNA complexes. The heterodimer supports stable fork pausing at the polar fork barriers of the *S.p.* mating type locus and the rDNA replication fork barriers (RFBs) of both *S.c*. and *S.p*. ^25–28^. At these programmed fork arrest sites Tof1/Swi1 ensures that adjacent sequences are replicated in a unidirectional manner – at the *S.p.* mating type locus this allows placement of the mating type “imprint”, and in the rDNA repeats, it minimises potentially deleterious head on collisions with highly active rRNA polymerases ^25–28^. This pausing activity is also observed at other high affinity protein-DNA complexes including centromeres and RNA pol III bound tRNA promoters ^29^. Although it is generally assumed that Tof1-Csm3 act to protect replication fork stability and prevent chromosome fragility at pausing sites, the importance of Tof1-Csm3 in preventing DNA damage in these contexts has not been analysed previously.

Tof1-Csm3 are also required to prevent replication stress due to accumulated DNA topological stress. Accumulation of DNA topological stress hinders DNA unwinding by the replicative helicase ^30, 31^. DNA topological stress accumulation is minimized both by DNA topoisomerase action and diffusion of topological stress through the DNA fibre from the point of generation ^32^. Sites of accumulation of DNA topological stress are generally associated with chromatin contexts that prevent diffusion of the stress along the chromatin fibre e.g., high affinity protein-DNA sites and nuclear envelope attachment regions ^33, 34^. Using ChIP-SEQ (<u>Ch</u>romatin Immuno<u>p</u>recipitation followed by next generation <u>seq</u>uencing) for H2AP/γH2AX, we have previously shown that the centromeres and the rDNA accumulate DNA damage during S phase when Top2 is depleted from cells, consistent with high affinity protein-DNA complexes and nuclear envelope association causing DNA topological stress accumulation. Tof1-Csm3 prevent replication stress due to accumulated DNA topological stress by recruiting eukaryotic topoisomerase I (Top1) to the replication fork through a direct interaction with the C terminus of Tof1 ^11, 35, 36^. The direct recruitment of Top1 to the replication fork likely rapidly resolves any accumulated DNA topological stress relieving associated replication stress. However, this prediction has not yet been directly tested.

Although not critical for bulk DNA synthesis, loss of *TOF1* or *Timeless* expression causes a marked increase in cellular DNA damage markers ^34, 37^. It is not known which of the different roles of Tof1/Timeless primarily prevent this increase in constitutive DNA damage. Potentially, the increased DNA damage could be associated with a generalised increase in fork uncoupling in cells lacking Tof1/Timeless and/or slow replication increasing the frequency of unreplicated regions persisting into mitosis. Alternatively, DNA damage could be due to a failure to pause replication at constitutive pausing sites (e.g. RNA pol III bound promoters) or caused by accumulated DNA topological stress causing fork arrest. In order to identify the primary causes of increased DNA damage in FPC-deficient cells, here we use genome wide assays in cells either lacking Tof1 or Mrc1 proteins or expressing mutant forms of Tof1 to identify the genomic contexts where Tof1 functions to protect *S.c.* cells from replication stress.

## RESULTS

### Deletion of Tof1 does not increase DNA damage at defined fork pausing sites and decrease DNA damage at centromeres and the rDNA

In *S.c.* the DNA damage sensing kinases Mec1/ATR and Tel1/ATM phosphorylate histone H2A at Serine 129 to generate H2AS129P (H2AP) ^38^. The equivalent action by ATR and ATM on the H2AX histone generates γH2AX in higher eukaryotes. The accumulation of H2AP in a chromosomal region occurs in response to either a DNA double strand break or exposure of ssDNA in that region ^39^. Loss of either yeast Tof1 or human Timeless leads to H2AP/γH2AX accumulation in cells ^34, 37^. In order to investigate where in the genome Tof1 prevents the local accumulation of DNA damage, we performed H2AP and H2A ChIP-SEQ in wildtype and *tof1Δ* exponentially growing cells. Previous ChIP on CHIP analysis of H2AP in wildtype cells observed that origins of replication (ARS sequences), centromeres, the upstream regions of tRNA and LTR sequences, telomeres, rDNA repeats and the silent mating type loci HML and HMR are all relatively enriched for DNA damage markers compared to other chromosomal regions ^3^. Using H2AP-H2A ChIP-SEQ, in wildtype cells we similarly observed strong enrichment of H2AP at telomeres, the rDNA repeats, the mating type loci HML and HMR and tRNA (enrichment at HML was partially obscured by high levels of enrichment in adjacent telomere proximal sequences) (Supplementary Figure 1 A-E). We also observed more modest enrichment of H2AP at origins of replication, centromeres and LTR transposons (Supplementary Figure 1 F-H). We also confirmed previous findings that repression of galactose inducible genes by growth in glucose increases local H2AP enrichment ^3^ (Supplementary Figure 2 A,B).

In *tof1*Δ cells we initially compared H2AP accumulation in regions where Tof1 is known to promote fork pausing; at the centromeres, tRNA and the RFB found in NTS2 of the rDNA repeats ^26, 27, 29^. Potentially loss of fork pausing at these replication stress inducing sites could lead to local DNA damage. However, in *tof1Δ* cells we observed either similar or lower levels of accumulations of H2AP at these sites. H2AP accumulated to *wt* levels at tRNA loci (Figure 1A) but was surprisingly reduced at the centromeres (Figure 1B) and across the rDNA repeat including the RFB region (Figure 1C). To ensure that loss of H2AP signal across the rDNA was not related to associated loss of rDNA copy number in *tof1Δ* cells (Supplementary Figure 2D), we normalized H2AP ChIP-SEQ signal in the rDNA to both H2A ChIP-SEQ (Figure 1C) and to input sequences (Supplementary Figure 2E). Both showed a loss of H2AP accumulation in *tof1Δ* cells across the rDNA. H2AP accumulation across other replication stress inducing contexts, including glucose repressed galactose inducible genes and origins of replication, were similar in both *wt* and *tof1Δ* cells (Supplementary Figure 2B, C).

**Figure 1.**
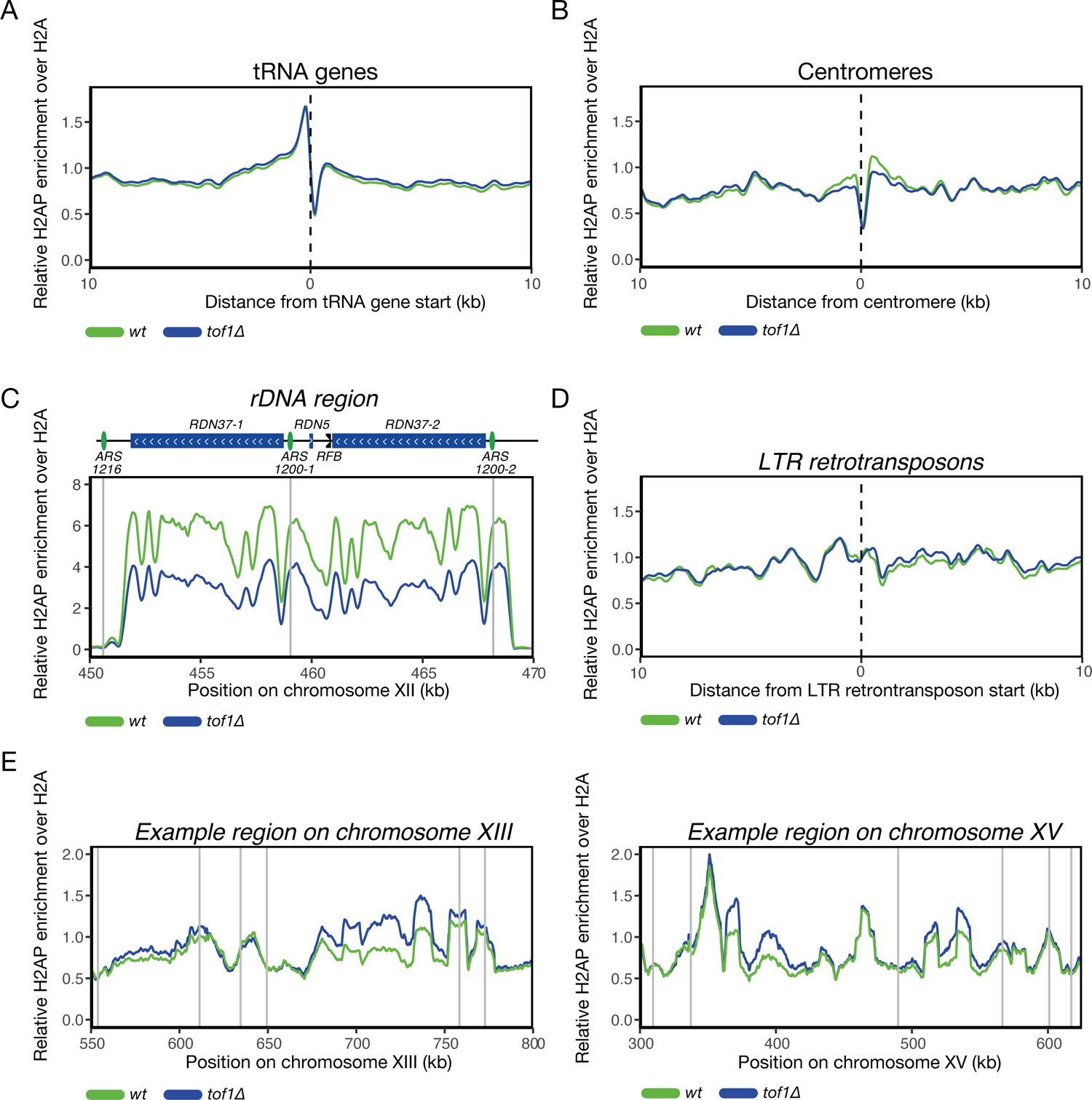
Deletion of Tof1 does not increase DNA damage at verified fork pausing sites and decreases DNA damage at centromeres and the rDNA Relative H2AP enrichment over H2A in *wt* and *tof1Δ* cells at (A) tRNA genes, (B) around centromeres, (C) at the rDNA region, (D) at LTR retrotransposons and (E) at two example regions on chromosome XIII (left) and XV (right). Grey vertical lines indicate positions of ARS sequences in the region. Smoothing with moving average over 7 bins (350bp) for (A-D) and 200 bins (10 kb) for (E) was applied.

Although we did not observe increased DNA damage at discrete replication stress associated loci, we did observe some chromosomal regions that accumulated H2AP in *tof1Δ* cells. We noted slightly increased H2AP upstream of LTR sequences in *tof1Δ* cells (Figure 1D). Interestingly, we also observed strong accumulation of H2AP in *tof1Δ* cells in regions across the genome where origins of replication (ARS) sequences were relatively distal from one another (Figure 1E).

### Tof1 dependent fork pausing and DNA damage do not correlate genome wide

Our initial investigation of candidate genomic loci suggested that Tof1-dependent fork pausing and Tof1-dependent inhibition of DNA damage were not connected. However, previous studies have only assayed Tof1-dependent pausing at discrete individual loci using two-dimensional gel electrophoresis. To determine if Tof1-dependent pausing occurs genome wide at these loci and also occurs at elevated frequency in regions where we observed DNA damage in *tof1Δ* cells, we compared the replication dynamics of *wt* and *tof1Δ* cells using the TrAEL-SEQ (<u>Tr</u>ansferase-<u>A</u>ctivated <u>E</u>nd <u>L</u>igation <u>seq</u>uencing) assay ^40^. The TrAEL-SEQ technique detects 3’ nascent DNA end exposure at reversed forks. Previous analysis of wild type cells has shown that TrAEL-SEQ signal is strongly elevated at constitutive replication pausing sites across the *S.c*. genome ^40^, including the RFB, centromeres and tRNA genes. TrAEL-SEQ detects the formation of 3’ ends on both nascent W and C strands and therefore provides information on any directional bias at pause sites.

Our TrAEL-SEQ analysis of wildtype cells confirmed bidirectional pausing at the tRNA and centromere (Figure 2A, B) and unidirectional arrest of forks at the RFB (Figure 2C) ^40^. In *tof1Δ* cells we observed loss of TrAEL-SEQ signals at tRNA (Figure 2A) and centromeres (Figure 2B) and a strong reduction in signal at the rDNA RFB as expected (Figure 2C). The loss of TrAEL-SEQ signal at the RFB was unidirectional, focused on forks proceeding from the proximal origin sequences (Figure 2C). Interestingly, loss of Tof1 also led to a unidirectional general decrease in TrAEL-SEQ signal across the rDNA in the same direction as bulk RNA transcription, suggesting that Tof1 supports transitory fork pausing during co-directional replication-transcription collisions (Figure 2D). At centromeres and tRNA, bi-directional loss of TrAEL-SEQ signal in *tof1Δ* cells was also observed, consistent with loss of pausing from forks approaching these structures from either direction (Figure 2A, B) ^40^. These findings are in accordance with the model that Tof1 supports fork pausing at a range of replication impeding structures throughout the genome.

**Figure 2.**
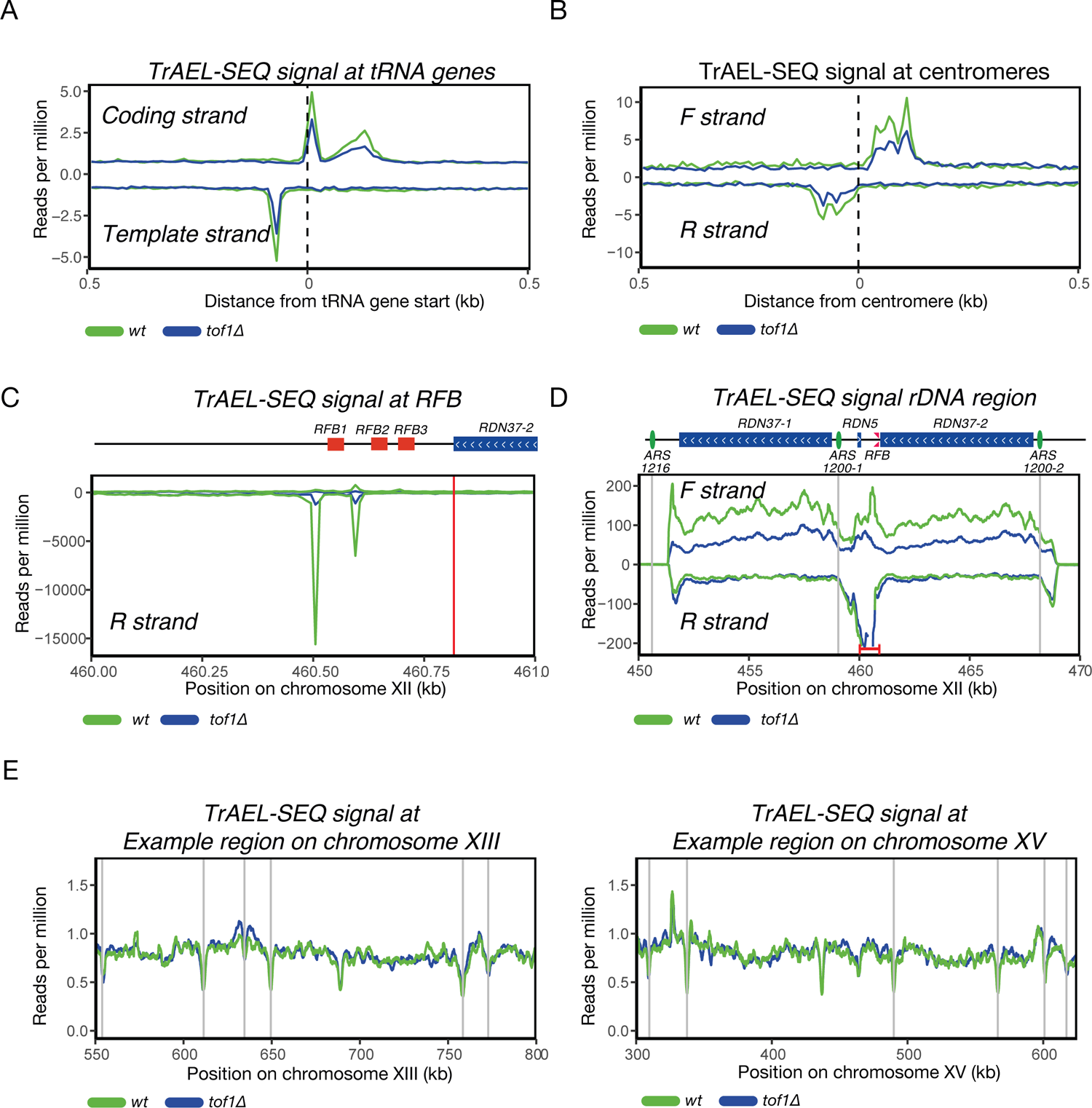
Tof1 dependent fork pausing and DNA damage do not correlate genome wide Directional TrAEL-SEQ signal at (A) tRNA genes, (B) centromeres, (C) rDNA replication fork barrier and (F) over the entire rDNA region. (E) Cumulative (forward and reverse strand) TrAEL-SEQ signal at two example regions on chromosome XII (left) and XV (right). Grey vertical lines indicate positions of ARS sequences in the region. For (A B and C) no moving average was applied (bin size 10 bp). For (D) moving average over 20 bins (200 bp) and for (E and F) moving average over 200 bins (2 kb) was applied. In (D) within the region bracketed in red reads exceed the y axis. This region is shown to a full scale in panel (C).

We next assessed TrAEL-SEQ signal across the regions where H2AP accumulated in *tof1Δ* cells to ascertain whether these regions contained previously unrecognised Tof1 dependent pause sites. Comparison of TrAEL-SEQ signal in *wt* and *tof1Δ* across the *tof1Δ* H2AP accumulating regions failed to show any substantiative decrease in TrAEL-SEQ signal in *tof1Δ* cells (Figure 2E). In summary we could not define any linkage between Tof1 dependent fork pausing and DNA damage accumulation.

### Tof1 protects long replicons from DNA replication stress

Visual inspection of the regions that did accumulate DNA damage in *tof1Δ* cells indicated a relative lack of ARS sequences in the vicinity of the H2AP signal. A regional absence of ARS sequences is associated with longer replicons. To test the hypothesis that replication stress in *tof1Δ* cells was preferentially occurring in longer replicons, we took all ARS sequences that have been assessed as likely to fire in most cell cycles (efficiency > 40 – based on ^41^) and used these sites to subdivide the genome into regions either likely replicated as part of a short replicon (20kb to 50kb) or a long replicon (>60kb) (only replicons within which other origins were relatively unlikely to fire (efficiency < 20) were considered) ^42^. We then assessed the average change in H2AP across different replicon sizes in *wt* and *tof1Δ* cells. Loss of Tof1 (*tof1Δ*) causes a marked increase in H2AP in long replicons while showing little effect in short replicons (Figure 3A).

**Figure 3.**
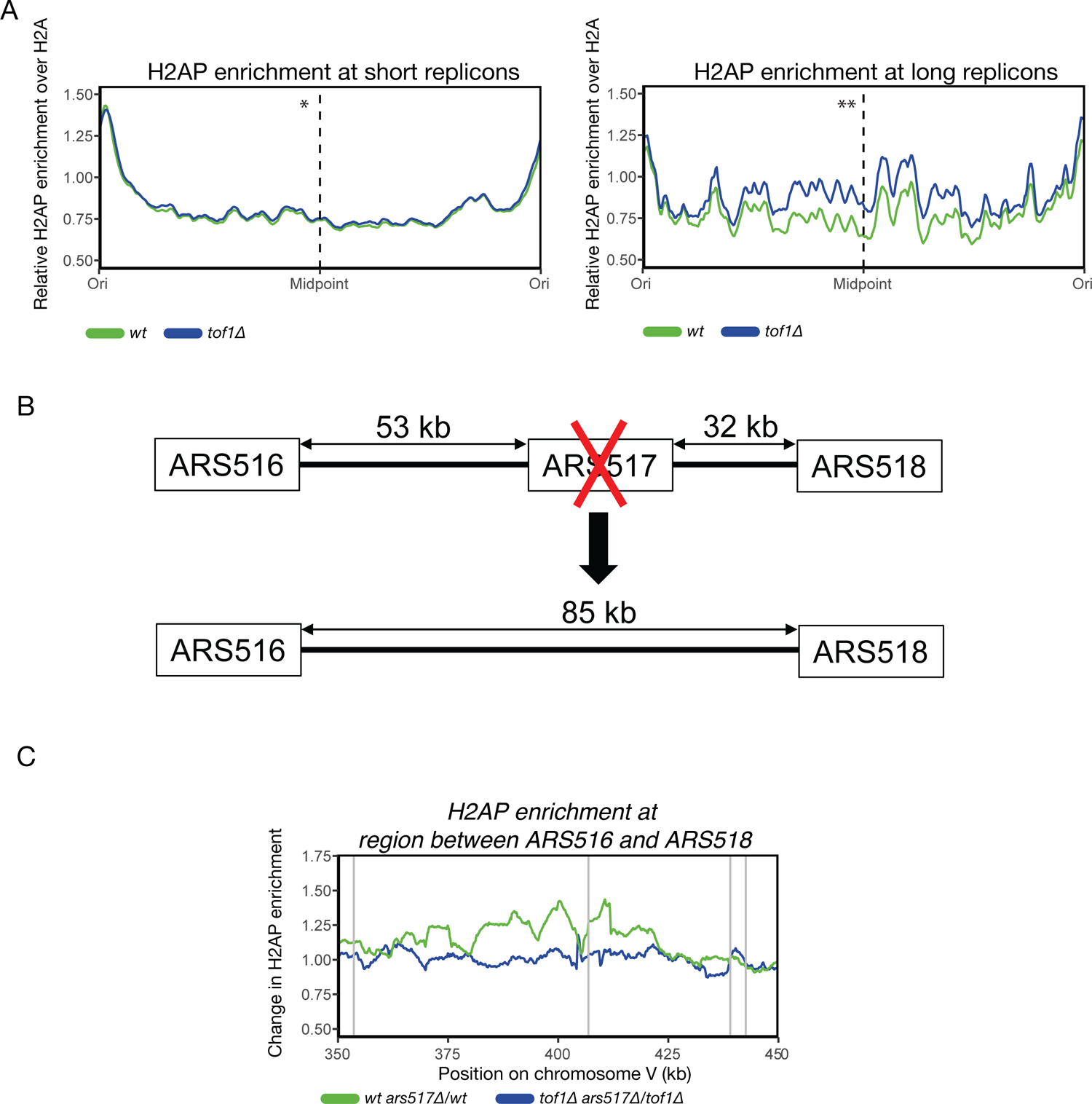
Tof1 protects long replicons from DNA replication stress (A) Relative H2AP enrichment in *wt* and *tof1Δ* cells over short replicons (20kb to 50kb) (left) and long replicons (>60kb) (right). * p-value of *tof1Δ* against *wt* in short replicons: = 0.8688. ** p-value of *tof1Δ* against *wt* in short replicons: = 0.004. (B) Schematic representation of the conversion of two relatively short replicons into one long replicon. (C) Ratio between relative H2AP enrichment in the presence or absence of ARS517 origin in *wt* and *tof1Δ* cells. Grey vertical lines indicate positions of ARS sequences in the region. Smoothing with moving average over 20 bins (1 kb) for (A) and 100 bins (5 kb) for (C) was applied.

To directly test the link between replicon length and DNA damage in *tof1Δ* cells, we converted two relatively short replicons that did not accumulate DNA damage in *tof1Δ* cells into one long replicon (Figure 3B). Our model predicts that the generation of a long replicon should specifically lead to accumulation of DNA damage in this region in *tof1Δ* cells but not in wildtype cells. We deleted the active early origin ARS517 to convert the two predicted replicons between ARS516 to ARS517 and ARS517 to ARS518 to one large replicon extending from ARS 516 to ARS 518 (Figure 3B). Comparing cells with and without ARS517 we observed that extension of replicon size had little effect on H2AP enrichment in wildtype cells (Figure 3C). In contrast, in *tof1Δ* cells the generation of an expanded replicon by deletion of ARS517 exhibited a marked accumulation of H2AP between ARS516 and ARS518 (Figure 3C). This shows that the accumulation of DNA damage in *tof1Δ* cells in longer replicons is not dependent solely on the underlying sequence. Rather, it demonstrates that the DNA damage in *tof1Δ* cells is a result of increased distance between origins of replication.

### Mrc1 protects long replicons from DNA replication stress

Tof1 and Mrc1 work together as part of the FPC complex to promote rapid and stable DNA replication ^26, 43–45^. Since we observed above that the DNA damage observed in *tof1Δ* cells is distinct from the fork pausing role of the FPC, which is Mrc1 independent, we sought to examine if DNA damage accumulation is linked to the interaction between Tof1 and Mrc1. Using H2AP/H2A ChIP-SEQ to examine *mrc1Δ* cells, we observed strong accumulation of H2AP in in the same long replicons affected in *tof1Δ* cells with only relatively minimal accumulation of H2AP in short replicons (Figure 4A). H2AP signal distribution in the affected regions of *mrc1Δ* was very similar to the profile of *tof1Δ* cells but with a notably higher intensity (Figure 4B). This indicates that loss of Mrc1 disrupts DNA replication in the same long replicons as Tof1, but in a more penetrative fashion.

**Figure 4.**
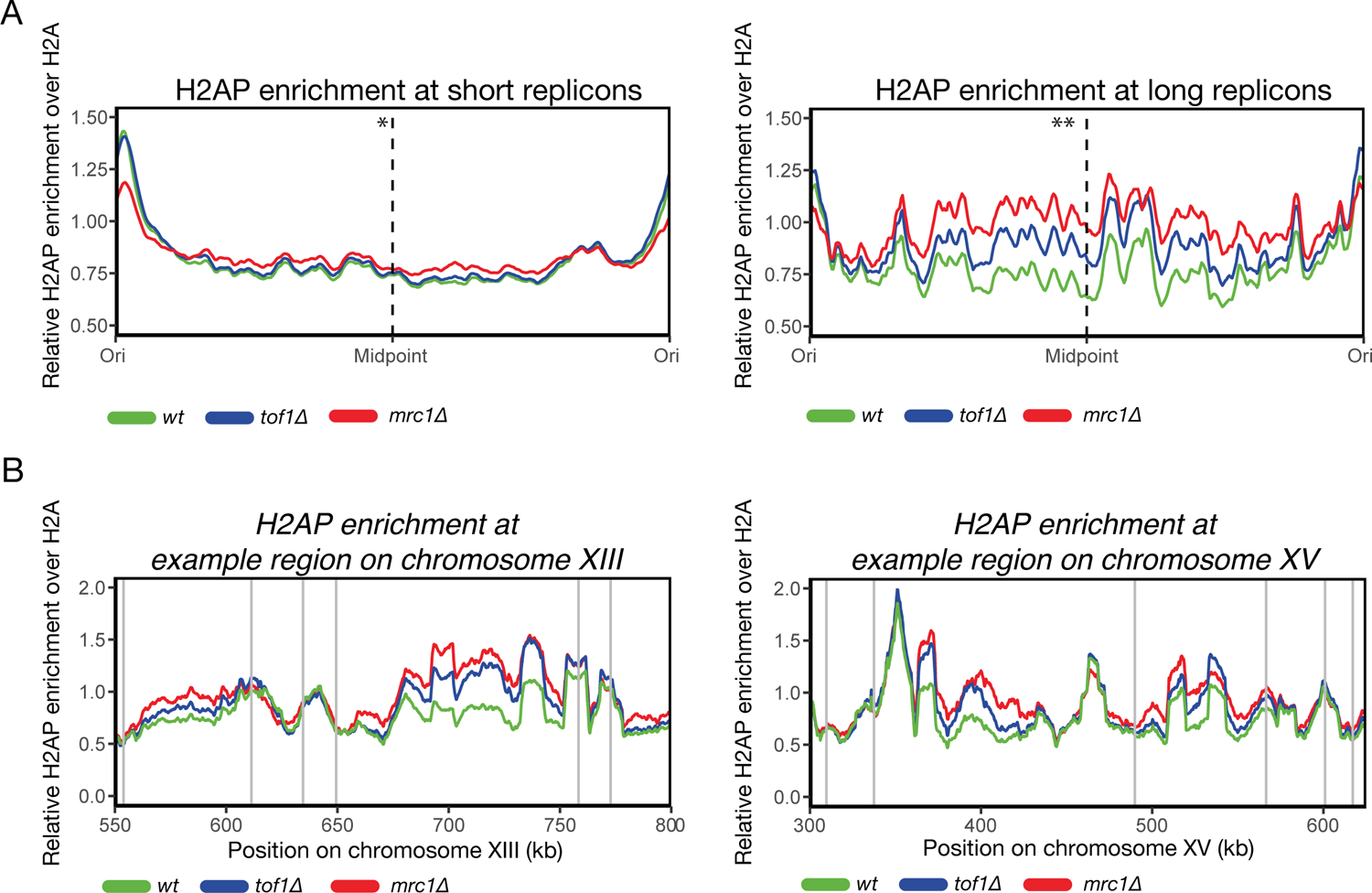
Mrc1 protects long replicons from DNA replication stress (A) Relative H2AP enrichment in *wt*, *tof1Δ* and *mrc1Δ* cells over short replicons (20kb to 50kb) (left) and long replicons (>60kb) (right). * p-value of *mrc1Δ* against *wt* in short replicons: = 0.534. ** p-value of *mrc1Δ* against *wt* in short replicons: = 6.226 x 10^-7^. (B) Relative H2AP enrichment in *wt*, *tof1Δ* and *mrc1Δ* cells at two example regions on chromosome XII (left) and XV (right). Grey vertical lines indicate positions of ARS sequences in the region. Smoothing with moving average over 20 bins (1 kb) for (A) and 200 (10 kb) for (B) was applied.

### Under-replication and persistent DNA damage occurs in long replicons in mrc1Δ cells

To further characterize the apparent difference in the intensity of constitutive DNA damage in *mrc1Δ* cells relative to *tof1Δ*, we examined both the cell cycle variation in DNA damage in *mrc1Δ* or *tof1Δ* cells and the relative propensity for under-replication of damaged regions of the genome in *mrc1Δ* or *tof1Δ* cells. Although H2AP accumulates in *tof1Δ* exponentially growing cells relative to wildtype, this was not observed in either G1 synchronized (treated with alpha factor) or mitotically arrested cells (arrested with nocodazole) (Figure 5A). This argues that H2AP accumulates primarily in S phase cells and that the associated DNA damage is not maintained in mitosis. In contrast, H2AP was highly elevated in *mrc1Δ* cells relative to wildtype in both exponential and mitotic post-replicative cells (arrested by nocodazole) while displaying similar levels damage to *wt* cells in G1 (Figure 5A). This indicates that DNA lesions generated during S phase in *mrc1Δ* cells either accumulate to a level where DNA repair kinetics are insufficient to ensure removal of the lesions before entry into mitosis, or the loss of the checkpoint signalling functions of Mrc1 result in delayed repair. Either scenario is consistent with the *RAD9* dependent DNA damage pathway being essential for survival in *mrc1Δ* cells ^22^. Loss of Mrc1 has also been reported to cause detectable under-replication of cells, as assayed by copy number variation^46^. To compare the relative states of under-replication in *mrc1Δ* and *tof1Δ* cells in short and long replicons relative to wildtype we used Next Generation Sequencing of exponentially growing cells to assess copy number variation in the two backgrounds ^47^. In *mrc1Δ* cells we detected under-replication in long but not short replicons relative to wildtype (Figure 5B). In *tof1Δ* cells we did not observe copy number differences in either short or long replicons relative to wildtype (Figure 5B). This suggests that the levels of replication stress and resultant under-replication and DNA damage are substantially higher following loss of Mrc1 relative to loss of Tof1.

**Figure 5.**
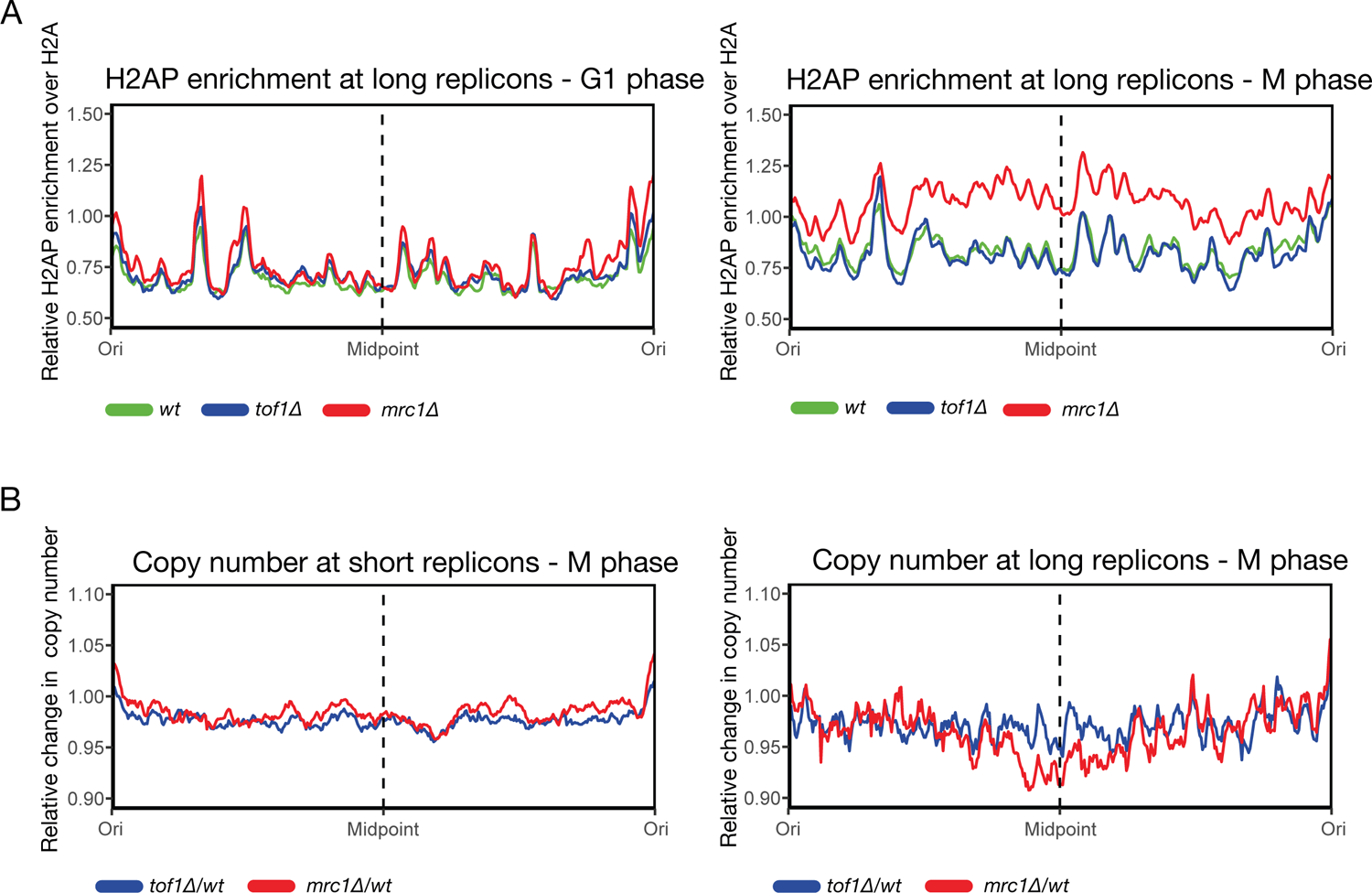
Under-replication and persistent DNA damage is focused in long replicons in mrc1Δ cells (A) Relative H2AP enrichment over H2A in *wt*, *tof1Δ* and *mrc1Δ* cells over long replicons (>60kb) in G1 synchronized (treated with alpha factor) (left) and mitotic cells (arrested with nocodazole) (right). (B) Ratio between *tof1Δ* or *mrc1Δ* over *wt* reads per million of input counts (relative change in copy number) across short replicons (20kb to 50kb) (left) and long replicons (>60kb) (right) in mitotic cells (arrested with nocodazole). Smoothing with moving average over 20 bins (1 kb) was applied. Correct cell cycle synchronisation of all cultures processed for (A) and (B) was confirmed by FACS for DNA content – Supplementary Figure 4.

### ssDNA accumulates in long replicons in mrc1 cells

Prior studies have shown that both Tof1 and Mrc1 are required to prevent uncoupling of helicase and polymerase activities at replication forks when cells are subjected to replication stess ^44^. Uncoupled regions are marked by increased exposure of ssDNA and the chromatin binding of the ssDNA binding protein RPA (which is composed of Rfa1, Rfa2 and Rfa3 in *S.c.*). To determine if increased exposure of ssDNA is a feature of the DNA damage that accumulates in long replicons in *mrc1Δ* and *tof1Δ* cells we performed Rfa1-ChIP in exponentially growing *wt*, *mrc1Δ* and *tof1Δ* cells. In *tof1Δ* cells Rfa1 ChIP-SEQ showed no significant accumulation of Rfa1 ChIP-SEQ signal across long replicons (Figure 6). In contrast, in *mrc1Δ* cells we observed strong accumulation of Rfa1 ChIP-SEQ signal, primarily in the mid regions of long replicons (Figure 6). In contrast, increased Rfa1 ChIP-SEQ signal was not observed in short replicons (Figure 6, left). Therefore, markers of fork uncoupling are only detectable in *mrc1Δ* cells and not *tof1Δ* cells. This data further supports the notion that the replication disruption in long replicons caused by loss of Mrc1 is quantitatively higher than loss of Tof1 function.

**Figure 6.**
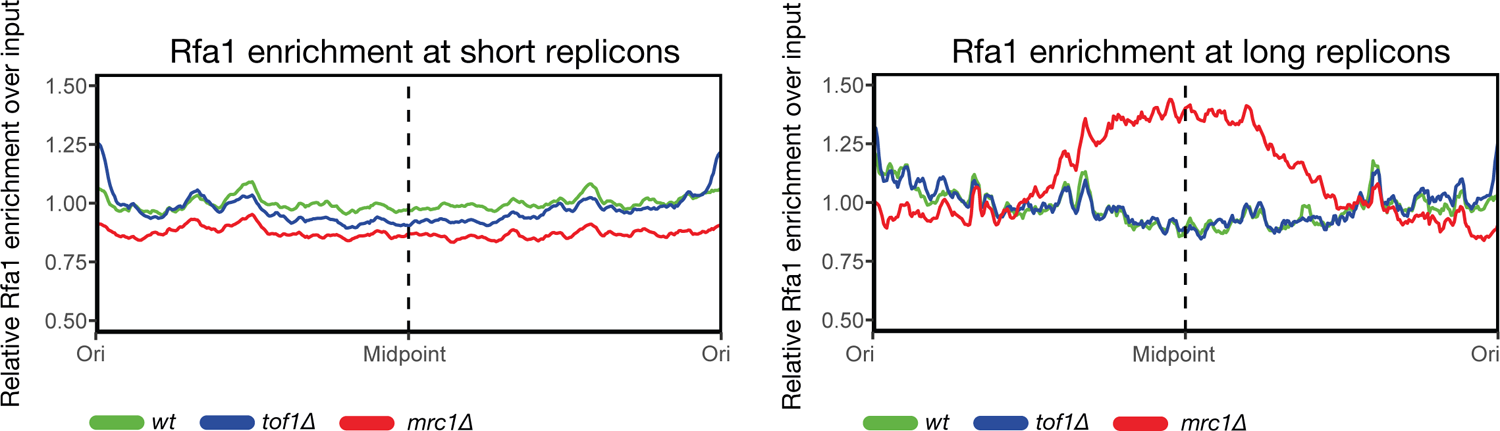
ssDNA accumulates in long replicons in mrc1 cells. Relative Rfa1 enrichment over input in wt, *tof1Δ* and *mrc1Δ* cells over short replicons (20kb to 50kb) (left) and long replicons (>60kb) (right) in exponential cells. Smoothing with moving average over 20 bins (1 kb) was applied.

### Tof1 recruits Top1 to suppress DNA damage accumulation at the centromeres and rDNA in rapidly replicating FPC+ cells

As part of the FPC, Mrc1 and Tof1 are required both for replication checkpoint signalling and, separately, for rapid and stable replication fork elongation ^20, 21, 26, 43, 44^. Previously we have characterised a truncation of Tof1, *tof1 627*, that maintains checkpoint signalling in response to hydroxyurea, but is defective for interaction with Csm3 and Csm3 linked functions ^36^. If replication checkpoint signalling was primarily required to prevent FPC linked DNA damage in long replicons we would predict that expression of *tof1 627* would suppress DNA damage accumulation due to loss of checkpoint signalling. However, cells expressing *tof1 627* showed very similar accumulations of H2AP to *tof1Δ* cells (Supplementary Figure 3). Therefore, restoration of replication checkpoint signalling did not detectably rescue DNA damage in long replicons, arguing that the damage is due to the loss of the rapid and stable replisome supported by all of the FPC factors including Csm3.

We next sought to test the importance of Tof1’s role in resolving DNA topological stress during DNA replication. Tof1 minimises replication stress caused by DNA topological stress by recruiting Top1 to the fork through a direct interaction in its flexible C terminal region ^34–36^. The *tof1 997* mutant is proficient in fork pausing and replication checkpoint activation but does not interact with Top1 and so is specifically defective for DNA topological stress resolution ^36^.To determine where Top1 recruitment prevents DNA damage in cells, we performed H2AP ChIP-SEQ in cells harbouring the *tof1 997* mutation. We did not observe any increase in H2AP accumulation in long replicons in this strain relative to *wt* (Figure 7A). Therefore, rapid DNA topological stress relaxation is not required for faithful replication of these regions. However, we did observe increased H2AP relative to H2A around centromeres and across the rDNA repeats in *tof1 997* cells (Figure 7B). Both centromeres and the rDNA accumulate DNA topological stress dependent DNA damage around centromeres and across the rDNA ^48^. Due to the propensity of these regions to accumulate topological stress we would predict that the loss of Top1 recruitment to the fork in *tof1 997* cells would exacerbate replication disruption. However, this model would appear to contradict our earlier observations that complete loss of Tof1 function caused a decrease in H2AP accumulation at both centromeres and across the rDNA (Figure 1B and C).

**Figure 7.**
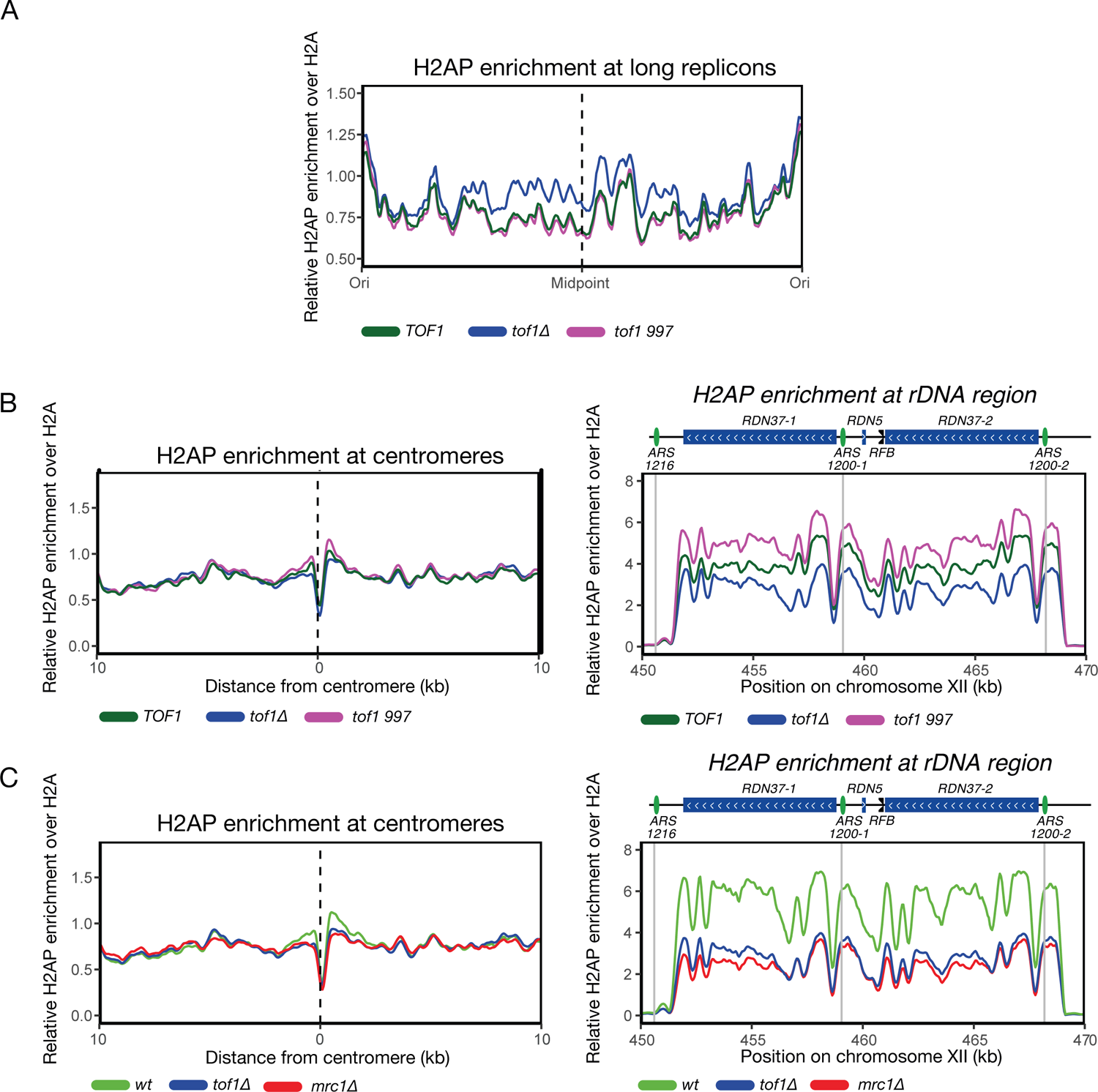
Tof1 recruits Top1 to suppress DNA damage accumulation at the centromeres and rDNA in rapidly replicating FPC+ cells (A) Relative H2AP enrichment in TOF1*wt*, *tof1Δ* and *tof1 997* cells over long replicons (>60kb) (B) Relative H2AP enrichment in TOF1*wt*, *tof1Δ* and *tof1 997* cells around centromeres (left) and rDNA (right). (C) Relative H2AP enrichment in wt, *tof1Δ* and *mrc1Δ* cells around centromeres (left) and rDNA (right)). Smoothing with moving average over 20 bins (1 kb) was applied for (A), and 7 bins (350 bp) for (B and C).

These findings can be reconciled if the rapid and stable replication provided by Tof1 (and Mrc1) is a cause of increased DNA topological stress in the centromeric and rDNA regions. In this model Tof1 dependent rapid unwinding by the helicase increases the frequency of generation of overwinding ahead of the fork. Increased overwinding then requires increased topoisomerase activity around the fork to prevent DNA topological stress accumulating to the extent that it stalls replication. The specific recruitment of Top1 to the fork by the Tof1 C-terminal region would promote increased topoisomerase activity around the fork. If rapid and stable replication was causing the high levels of DNA topological stress resolution, loss of Mrc1, like *tof1Δ* should cause a reduction in DNA damage at centromeres and across the rDNA. We found that loss of Mrc1 does reduce the levels of H2AP across both centromeres and the rDNA relative to wildtype demonstrating that increased constitutive DNA damage in these regions is due to an Mrc1-Tof1 linked function (Figure 7C). Therefore, combining all our observations, we conclude that Tof1-Mrc1 are required to promote stable and rapid DNA replication to ensure the faithful duplication of long replicons (Figure 8A, B). But stable and rapid replication comes at the expense of increased DNA topological stress ahead of the fork (Figure 8C). Thus, rapid replication causes high levels of DNA topological stress ahead of the fork. Although this is tolerated across most of the genome, it which makes cells acutely sensitive to loss of topoisomerase activity in regions prone to topological stress accumulation (Figure 8C), such as the centromeres and the rDNA of budding yeast ^48^.

**Figure 8.**
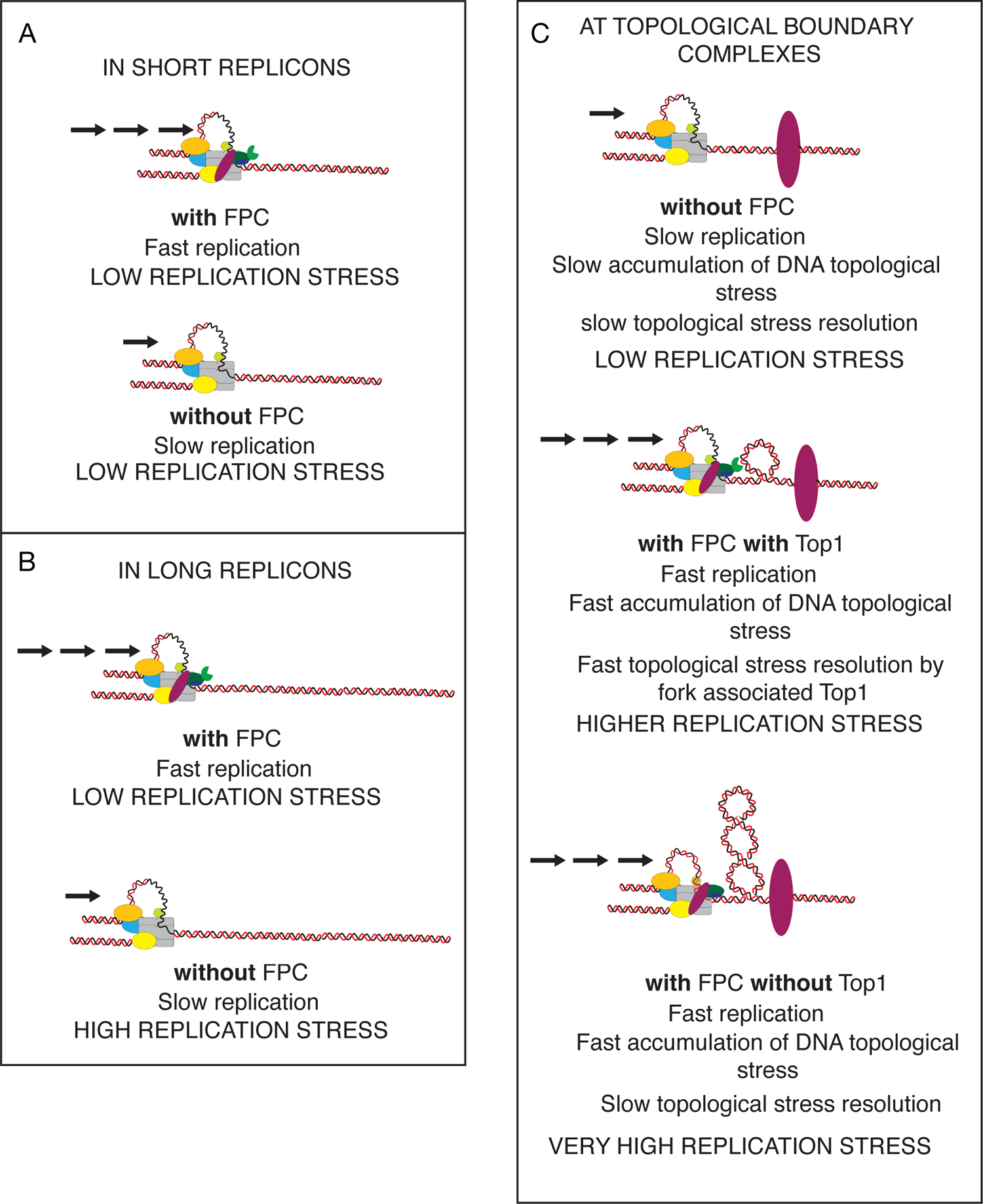
Model of how the FPC prevents replication stress at long replicons but also promotes DNA topological stress at the CEN and rDNA With the FPC intact the replisome is capable of rapid and stable replication (A). Without the FPC, DNA replication is relatively slow. Across short replicons both rapid and slow replication are sufficient to ensure completion of replication without accumulating significant DNA damage. In contrast across long replicons (B) rapid and stable replication is required for full completion of the long replicon. Slow replication leads to frequent failure of replication before completion. In regions containing DNA topological boundary complexes (C) the DNA topological stress generated by DNA unwinding cannot diffuse away from the site of generation and will accumulate locally, potentially stalling replication. In the absence of an intact FPC, DNA unwinding is slow and therefore DNA topological stress accumulation is slow. Therefore, diffusing topoisomerases are sufficient to prevent stress accumulating to the point where it could stall replication (top panel). However, if the FPC is intact replication is fast, DNA unwinding is fast and therefore DNA topological stress accumulates quickly. In this situation topoisomerase I needs to be actively attracted to the replication fork by protein-protein interaction to prevent DNA topological stress accumulating to very high levels (middle panel). Without active topoisomerase recruitment the stress accumulates to a level where replication stalling occurs frequently (bottom panel).

## DISCUSSION

DNA replication stress is a recognized hallmark of pre-oncogenic cells and likely a precondition of progression into the fully cancerous state ^49^. The FPC proteins Timeless and Claspin become overexpressed as cells progress from the precancerous to cancerous state ^50^, indicating that their functions are required to maintain cellular viability in the face of increased oncogene induced replication stress. Here, we show that the yeast homologues of Timeless and Claspin in the FPC, Tof1 and Mrc1, are required to prevent DNA damage accumulating in long replicons (Figure 8 A,B). We also find that the rapidly replicating replisome generated by the FPC increases DNA damage in regions prone to DNA topological stress, necessitating the active recruitment of Top1 to the fork to minimise damage in these zones (Figure 8C).

Long replicons are amongst the most frequently damaged genomic regions in cancerous cells ^49^, indicating that they present specific challenges for DNA replication. Since Tof1/Timeless and Mrc1/Claspin are established regulators of replication speed and fork coupling ^26, 43, 51^ our findings show that by potentiating rapid and stable replication in long replicons these factors ensure the full and faithful duplication of the genome in each cell cycle (Figure 8 left). Although loss of either Tof1 or Mrc1 results in DNA damage in long replicons, they do not appear to be equally important in preventing DNA damage at these loci. Loss of Tof1 leads to an S phase focused increase in H2AP whereas loss of Mrc1 leads to a higher concentration of H2AP in long replicons which persists into M phase, whilst also causing extensive exposure of ssDNA formation in these regions. *In vitro* omission of Mrc1 from replication reactions causes greater loss in replication speed than omission of Tof1-Csm3 alone ^43^. Furthermore, although Tof1-Csm3 are required for strong association of Mrc1 with the replisome, Mrc1 is not required for the interaction of Tof1-Csm3 with the replisome ^52^. These observations argue that the primary role of Tof1-Csm3 in coping with replication stress is to stabilize and correctly orient Mrc1 in a manner that promotes stable, rapid DNA replication and prevents uncoupling and extensive exposure of ssDNA in long replicons. We cannot discount the possibility that the additional DNA damage and ssDNA exposure caused in *mrc1* mutants relative to *tof1* could be due to checkpoint related functions of Mrc1, for example through Mrc1 preventing the resection of stalled forks ^53^. At present the nature of the DNA lesions generated in long replicons following loss of the FPC is unclear. Increased frequency of uncoupling of helicase-polymerases above a certain period without FPC activity would seem the simplest pathway. However, we cannot exclude other possibilities such as slow replication causing the firing of low-frequency stochastic origins in longer replicons ^54^ that could be inherently more unstable in the absence of FPC activity.

Independently of its interaction with Mrc1, Tof1 promotes fork pausing at a range of replication stress inducing sites. We sought here to investigate if Tof1 promoted fork pausing was important for the accumulation of DNA damage at these sites in unchallenged cells. We found that loss of Tof1 function and fork pausing does not lead to increased levels of DNA damage at tRNAs, centromeres or the rDNA RFB despite our findings via TrAEL-SEQ that Tof1 promotes fork pausing at these sites on all chromosomes. This raises the questions as to why does Tof1 activity enforce fork pausing at these loci if not to prevent DNA damage? It has been argued that Tof1-Csm3 enforces pausing by impeding Rrm3 “sweepase” function, potentially by enforcing a specific replisome structure ^28^. Although Rrm3 function is not necessary for Tof1 dependent pausing ^35^, the Tof1-Csm3 heterodimer does enforce a specific replisome structure that optimizes Mrc1 functions and rapid replication ^7^. We postulate that pausing is simply a consequence of Tof1-Csm3 stabilising a replisome conformation that is inefficient at bypassing stable protein-DNA complexes.

Although Tof1 dependent fork pausing does not prevent DNA damage at fork pausing structures, we do find that loss of FPC activity influences genome stability around structures that accumulate DNA topological stress. Our data show that loss of Tof1 or Mrc1 reduces DNA damage accumulation at centromeres and across the rDNA repeats. We have previously reported that centromeres and the rDNA repeats are particularly susceptible to DNA topological stress related DNA damage ^48^. Therefore, the same activities that promote genome stability in long replicons, also contribute to replication dependent DNA damage across topologically stressed zones (Figure 8 A, B vs C). Replisomes containing Tof1-Mrc1 that can replicate rapidly and stably will generate particularly high levels of DNA topological stress ahead of the fork. In regions where topological stress diffusion is limited, this rapid accumulation of overwinding makes it imperative that topoisomerase action resolves the stress before it reaches critical, replication stalling, levels (Figure 8C). Centromeres and the rDNA are especially vulnerable to DNA topological stress accumulation due to high levels of cohesin in these regions ^48^. This combination explains why Mrc1-Tof1 activity cause additional replication problems in specific genomic contexts and provides an evolutionary imperative for Tof1 recruiting Top1 to the fork to counteract the potentially deleterious effects of rapid replication (Figure 8C).

This model indicates that while FPC functions are very important for the complete replication of the genome (and likely essential in some human cell types ^55^) their combined functions can be deleterious to genome stability in some chromosomal contexts. Such a model provides a rationale as to why both yeast Mrc1 and mammalian Timeless are targets of stress response mechanisms aimed at inhibiting maximal DNA replication dynamics in cellular contexts where rapid DNA replication could induce genome instability ^56, 57^. These findings highlight that modes of replication regulation beneficial in some contexts are deleterious in others, demonstrating that numerous contrasting pathways of replisome regulation are required in eukaryotes to balance the differing chromosomal and cellular challenges to faithful chromosome replication.

## ACKNOWLEDGMENTS

We thank Antony Carr, Ulrich Rass, Nicola Minchell and Rose Westhorpe for critical reading of this manuscript. This work was funded by the Biotechnology and Biological Sciences Research Council, United Kingdom, (BBSRC UK) Grants ref BB/N007344/1 and BB/S001425/1 (A.K., J.B.) and the Wellcome Trust (A.W., J.H.).

## AUTHOR CONTRIBUTIONS

Conceptualization, J.B.; Methodology, A.K., A.W. and J.H.; Investigation, A.K., A.W..; Software, A.K.; Formal Analysis, A.K.; Writing – Original Draft, J.B. A.K..; Writing – Review & Editing, J.B., A.K., A.W. and J.H.; Funding Acquisition, J.B. and J.H.; Supervision, J.B. and J.H.

## SUPPLEMENTARY FIGURE LEGENDS

### Materials and methods

#### Yeast strains

Strains were generated in the W303 background *(ade2–1 ura3–1 his3–11, trp1–1 leu2–3, can1–100*) and are listed in the Table 1.

#### Media and Cell Cycle Synchronization

For exponential ChIP-SEQ experiments in glucose, cells were grown at 25 °C in YP media with 40 mg/l adenine + 2% glucose to mid-log phase (∼10^7^ cells/ml).

For exponential ChIP-SEQ experiments in galactose, cells were grown to ∼0.7 x 10^7^ cells/ml at 25 °C in YP media with 40 mg/l adenine + 2% raffinose first, then 2% galactose was added, and cells were further incubated to reach ∼10^7^ cells/ml before collection.

For exponential experiments in glucose for TrAEL-SEQ, cells were grown at 30 °C in YP media with 40 mg/l adenine + 2% glucose to mid-log phase (∼10^7^ cells/ml).

Cell synchronizations were performed as described previously ^34^. Cultures were grown in YP media with 40 ml/l adenine + 2% raffinose to midlog phase, then 10 μg/ml alpha factor (Genscript) was added to arrest cells in G1 phase. After ∼120 min, when 90% of cells were unbudded (G1 phase) 2% of galactose was added. 20 minutes later 50 μg/ml doxycycline (Sigma-Aldrich) and an additional 5 μg/ml alpha factor was added. After a further 10 minutes the temperature was shifted to 37°C for 1h. Cells were released from G1-block by 3x wash with YP + 40 mg/l adenine + 2% raffinose + 2% galactose 50 μg/ml doxycycline followed by resuspension in the same media. For mitotic arrested samples, nocodozole (Sigma-Aldrich) was added to cultures at 10 μg/ml 45 minutes after time 0 min (addition of the first wash), and cells were collected at 95 min. For G1 arrested samples, 10 μg/ml alpha factor was added after 70 min and cells were collected at 160 min. Cell cycle phases were confirmed by Flow cytometry analysis (FACS)(Supp. Figure 4).

#### Flow cytometry analysis (FACS)

FACS was performed as described in ^34^. 500 μl of culture was collected by centrifugation and fixed with 70% ethanol. Cells were then RNAse treated in 1 ml of 50 mM Tris-HCl pH8 with 5 mg/ml RNaseA (Sigma-Aldrich) at 37°C overnight, then protease treated in 1 ml 5 mg/ml pepsin (Sigma-Aldrich) and 5 μl/ml concentrated HCl at 37°C for 30 minutes. Cells were then washed in 50 mM Tris-HCl pH8 and resuspended in 50 mM Tris-HCl pH8 with 0.5 mg/ml Propidium iodide (Sigma-Aldrich). After brief sonication FACS was performed using BD Accuri C6 sampler. Analysis was carried out using FCS express 4 flow software. Data for FACS analysis is shown in Figure S4.

#### ChIP-SEQ

ChIP-SEQ experiments were performed as described previously ^48^. 25 ml of cultures were used per each antibody used for ChIP. Cells were washed and resuspended in YP media, then 1% formaldehyde (Sigma-Aldrich) was added, and cells were incubated for 45 min at 25°C. To quench the formaldehyde 125 mM glycine (Alfa Aesar) was added followed by 5 min incubation. Cells were then washed with cold PBS, pelleted and frozen in liquid nitrogen. For a subset of experiments (see Supplementary Table 1.) normalization by adding *Schizosaccharomyces pombe* (*S. pombe*) cells containing the HO-endonuclease inducible system (AW507) were used (as described in ^48^). Briefly, primary cultures of AW507 were grown in EMM media supplemented with leucine (100 μg/ml) at 30°C overnight, then kept at logarithmic phase for a day, before diluting to a secondary culture to reach logarithmic phase (5×10^6^ cells) again the next day. Cells were then pelleted and re-suspended in pre-warmed EMM supplemented with leucine, histidine and uracil (100 μg/ml each) to induce HO endonuclease (Purg1loxON). After two hours of incubation at 30°C cells were fixed and collected as described above for *S. cerevisiae* cells.

Pellets from liquid nitrogen were resuspended in 500 μl SDS buffer (1% SDS, 10 mM EDTA, 5M Tris HCl, cOmplete Tablets, Mini EDTA-free EASYpack (Roche), PhosSTOP (Roche)), then lysed in a FASTPREP machine, 5 rounds of 1 min at 6.5 power, with 200 μl of 0.5 mm silica beads. Silica beards were then separated and discarded and IP buffer (0.1% SDS, 1.1% Triton-X-100, 1.2 mM EDTA, 16.7 mM TRIS HCl (pH8), cOmplete Tablets, Mini EDTA-free EASYpack (Roche), PhosSTOP (Roche)) was added to the lysate to a final volume of 1 ml. Focused-Ultrasonicator (Covaris, M220) was then used to sonicate the samples (Average incident power – 7.5 Watts, Peak Incident Power – 75

Watts, Duty Factor – 10 %, Cycles/Burst – 200, Duration – 20 min). For 50 ml starting culture the supernatant was diluted to 5 ml in total. 50-50 μl of protein A Dynabeads (Invitrogen) and protein G Dynabeads (Invitrogen) were mixed and washed 3 times in IP buffer then added to the samples and was incubated for 2 h at 4°C. After discarding the beads 2-2 ml of the supernatant was incubated with either H2A 1:500 (active motif) or 1.6 μg/ml H2AP (Abcam) on a rotating wheel at 4°C for 15 – 20 h. The rest of the sample was kept for input. If RFA1 ChIP-SEQ was also performed, the volume of the starting culture was increased to 75 ml and the sample after sonication was diluted to 7.5 ml in total and 75-75 μl of protein A and G Dynabeads were used to reduce unspecific binding and RFA1 antibody (1:10000, Agrisera) was added to 2 ml of the pre-cleaned sample followed by incubation on a rotating wheel at 4°C for 15 – 20 h. 30-30 μl of protein A and G Dynabeads were washed 3x with IP buffer and was added to each antibody reaction and incubated at 4°C for 4 h.

Beads were then washed at 4°C for 6 min in TSE-150 (1% Triton-X-100, 0.1% SDS, 2 mM EDTA, 20 mM Tris HCl (pH8), 150 mM NaCl), followed by TSE-500 (1% Triton-X-100, 0.1% SDS, 2 mM EDTA, 20 mM Tris HCl (pH8), 500 mM NaCl), followed by LiCl wash (0.25 M LiCl, 1% NP-40, 1% dioxycholate, 1 mM EDTA, 10 mM Tris HCl (pH8)) and finally Tris-EDTA (TE pH8). Samples were then eluted in 400 μl elution buffer, for 30 min at room temperature. Input samples were prepared from 50 μl starting material mixed with 150 μl of elution buffer (200 μl final volume). For reverse crosslinking and protease treatment NaCl (500 mM final concentration) and proteinase K (Invitrogen, 500 μg/ml final concentration) was added and incubated at 65°C overnight. RNase treatment was then performed by adding DNase-free RNase (Roche, 25 μg/ml final concentration) followed by incubation at 37°C for 30 min. DNA was then purified with Qiagen PCR purification kit.

NGS libraries were prepared using NEBnext Ultra II library kit (NEB) with 13 cycles used for PCR enrichment. AMPure XP beads were used for size selection and PCR purification. DNA yield was measured by Qubit 2.0 Fluorometer (Life technologies). For input and RFA1 libraries 34 μl from the RFA1 IP samples and 1 ng DNA in 34 ul water from the input was used as starting material. To generate complementary strand for the ssDNA first 5 μl 10 x NEB2.1 buffer and 5 μl of random primers (8N, 3 mg/ml stock) were added and the samples were boiled at 95°C for 5 minutes and immediately placed to ice for 5 minutes. 5 μl 10 x dNTP with dUTP instead of dTTP (2 mM each) and 1 μl T4 polymerase (NEB) were then added, and the mixture was incubated at 37°C in a thermal cycler for 20 min. 5 μl 0.5 M EDTA (pH 8) was immediately added to stop the reaction. This was then used to generate libraries using Ultra II library kit (NEB) with 16 cycles for PCR enrichment. Paired end sequencing was performed using NextSeq 500 (42 bp reads from each side) system. For experiments with normalization using *S. Pombe* cells, aliquots (∼2.5 x 10^9^ cells) of *S. pombe* cell pellets were resuspended in 250 μl SDS buffer, and 1/1000 volume of the original *S. cerevisiae* culture (corresponding to 1:10 *S. pombe* to *S. cerevisiae* ratio) was added to each *S. cerevisiae* samples which were then processed the same way as described above.

#### ChIP-SEQ data analysis

Data analysis for ChIP-SEQ was performed as described previously ^48^. Illumina basespace (https://basespace.illumina.com/home/index) was used to generate FASTQ files from the sequencing reactions. H2A and H2AP sequences were aligned without trimming to a reference genome (R64-1-1, *Saccharomyces cerevisiae* S288c assembly from Saccharomyces Genome Database) using Bowtie 2 (http://bowtie bio.sourceforge.net/bowtie2/index.shtml). RFA1 reads were aligned to the same reference genome but the LTR-retrotransposons were masked.

Command for bowtie2 alignment:

`bowtie2 -p 14 -x [path to index folder] --trim3 0 --trim5 0 - 1 [Path and name of R1 fastq file] - 2 [Path and name of R2 fastq file] -S [name of the resulting .sam file]`

SAM files were then converted into sorted BAM files by using SAMtools (http://samtools.sourceforge.net/):

`samtools sort [name of the .sam file generated with bowtie2] - o [name for the resulting .bam file] -O bam -T [name for temporary file (optional, used if parallel nodes are used)]`

For RFA1 analysis duplicates were then removed using picard (https://broadinstitute.github.io/picard):

java -jar ∼/picard/picard-tools-1.138/picard.jar MarkDuplicates I= [name for the resulting .bam file] O=[name for the resulting without repeats.bam file] M= [name of metrix file.txt] REMOVE_DUPLICATES=true

BAM files were used for Model-based Analysis of ChIP-SEQ (MACS2). We used the ‘call peak’ function which also generates genome wide score data. These were used to generate fold enrichment tracks. Example command:

`macs2 callpeak -t [sorted BAM file from yh2a data]-c [sorted BAM file from h2a data]-f BAMPE -g 12100000 -n [name for output file] -B -q 0.01 –SPMR`

The data then was sorted into 50 bp bins, normalized to have a mean value of 1, smoothed moving averages indicated at each figure and used for meta data analysis and plotting using custom made R programs.

#### Relative copy number determination

Libraries for relative copy number determination were prepared as described for the input preparation for RFA1-ChIP. Reads were aligned LTR-retrotransposon masked reference genome, duplicates were removed using picard, and reads were summed to 50 bp bins using sam- to bincount program (https://github.com/yasukasu/sam-to-bincount) described in60: perl filepath/pe-sam-to-bincount.pl –i [name of the .sam file generated with bowtie2] –-trim5 0 –-strand –-end 0 –n 0 –w 50 –ref [Path and name of reference fasta file] Read per million values were calculated (rDNA values ignored) and values from forward and reverse strands were summed using custom R scripts.

#### TrAEL-SEQ

TrAEL-SEQ experiments were performed as described earlier in ^40^.

#### TrAEL-SEQ data analysis

UMI deduplicated mapped reads from TrAEL-SEQ experiments were generated as described in ^40^ . Mapped reads were then analysed using SeqMonk v1.47 (https://www.bioinformatics.babraham.ac.uk/projects/seqmonk/). Minimum mapping quality of 1 was applied, and reads were truncated to 1 nucleotide at the 5’ end. Running windows of probe size 10 bp and step size 10 bp were generated and the reads were exported to bedgraph file. Custom made R programs were then used to calculate reads per million values (reads around rDNA were ignored). Reads per million values were then smoothed by moving averages indicated at each figure for plotting using custom made R programs. When plotting metadata CUP1 region (+-5kb) was ignored.

## Data and Code Availability

Processed sequencing data were deposited in GEOxxxx

---

**Supplementary Figure 1.**
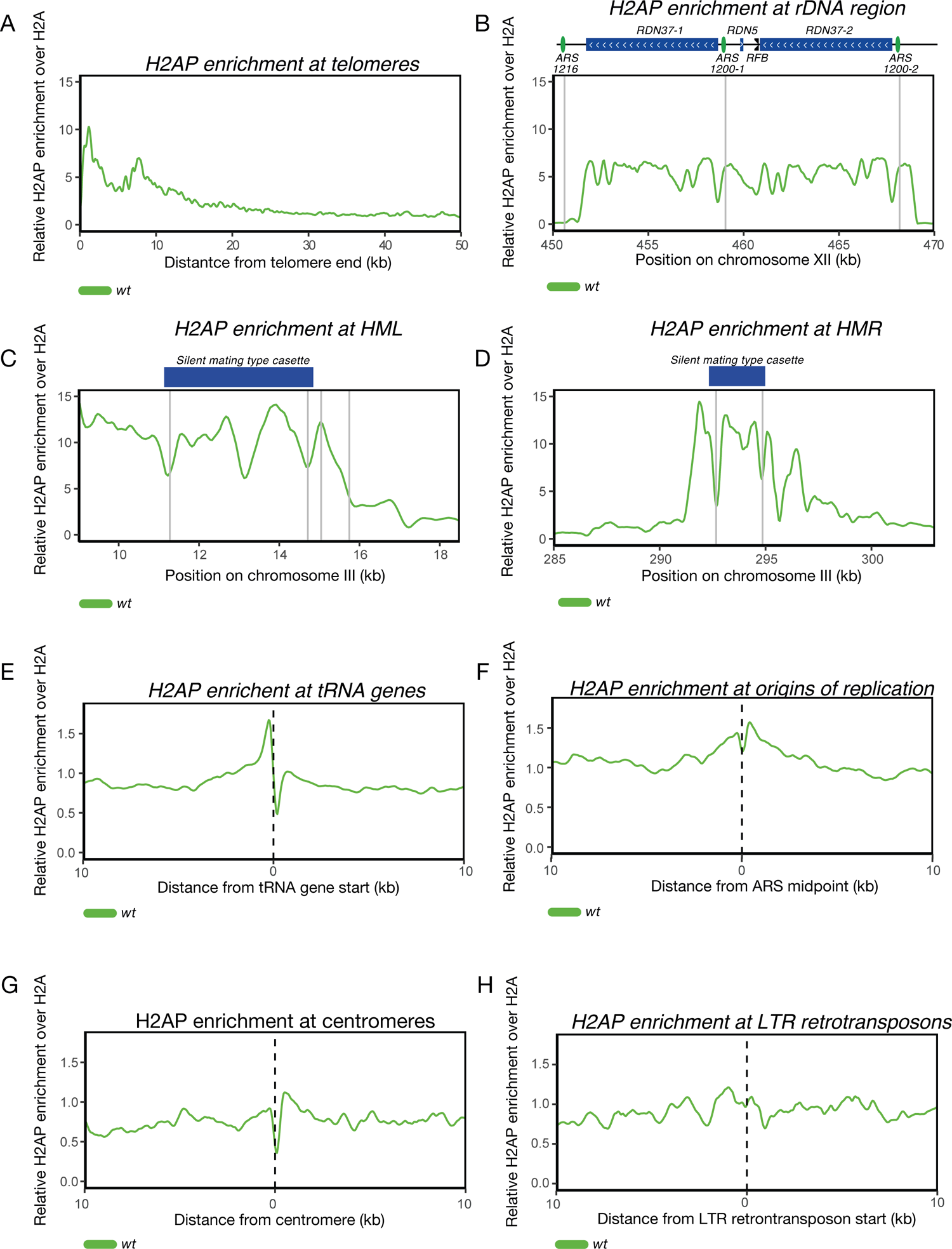
H2AP enrichment signal is increased in wt cells at specific regions Relative H2AP enrichment in wildtype cells at (A) telomeres, (B) rDNA repeats, (C) HML - silent mating type locus, (D) HMR - silent mating type locus (grey vertical lines indicate positions of ARS sequences in the HML and HMR regions), (E) upstream and downstream regions of tRNA, (F) origins of replication (ARS sequences), (G) centromeres and (H) LTR sequences. Smoothing with moving average over 7 bins (350 bp) was applied.

**Supplementary Figure 2.**
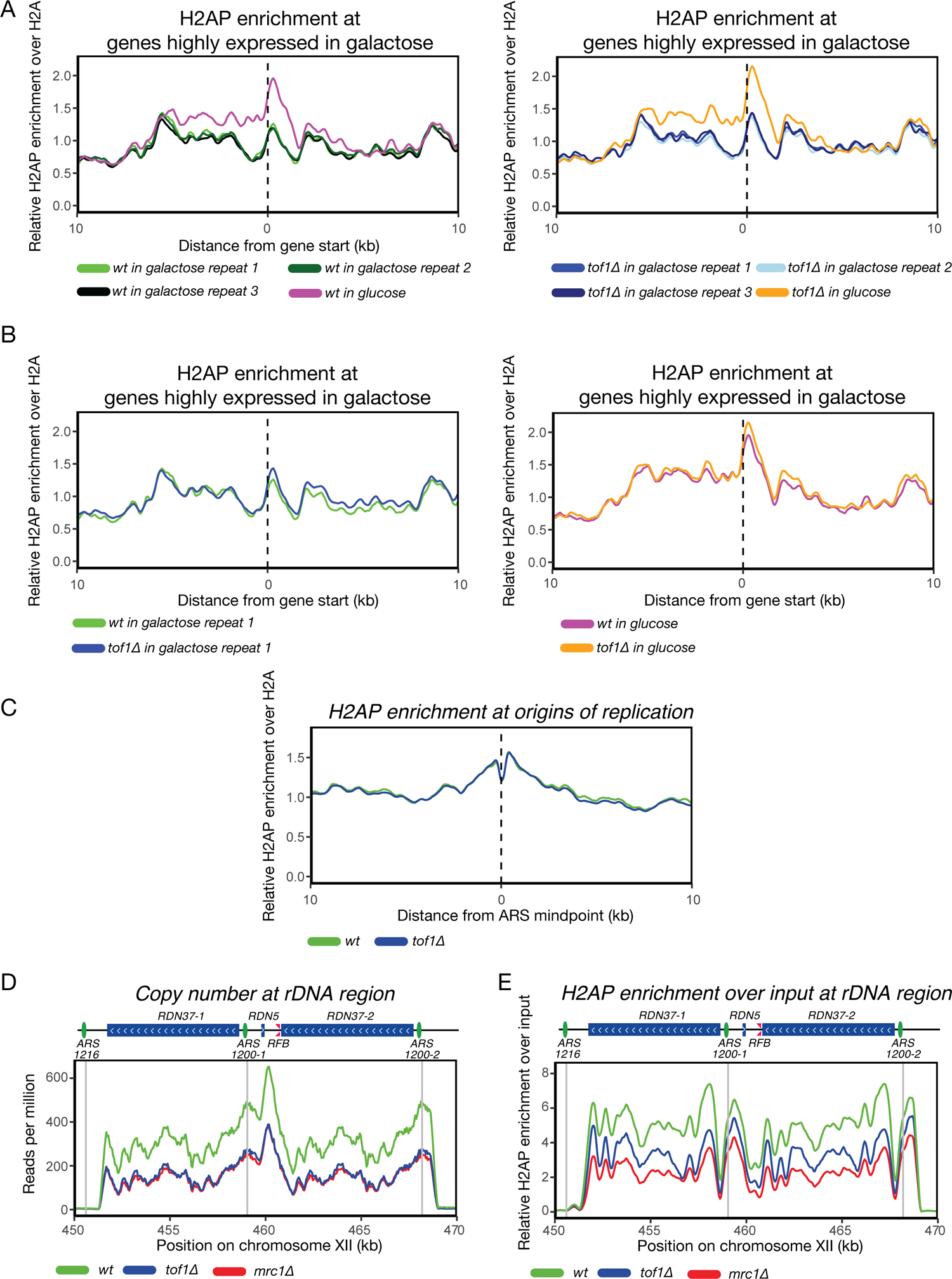
Repression of galactose inducible gene in glucose media causes H2AP accumulation both in wt and TOF1 deleted cells. H2A enrichment at origins of replications is not affected by loss of Tof1. H2AP enrichment at rDNA is not dependent on copy number Determination of highly expressed genes in galactose was based on the dataset in ^58^ expression in YP galactose vs reference pool >1.5 (A) Relative H2AP enrichment over H2A at genes highly expressed in galactose in wt (left) and tof1Δ (right) grown in media containing galactose or glucose. (B) Relative H2AP enrichment over H2A at genes highly expressed in galactose in wt and tof1Δ cells grown in media containing galactose (left) or glucose (right). (C) Relative H2AP enrichment over H2A around origins of replication in wt and in tof1 deleted cells. (D) Copy number across rDNA region. (E) The relative enrichment of H2AS129P over input ChIP across the rDNA repeats in wt and tof1Δ cells. Grey vertical lines indicate positions of ARS sequences in the region. Smoothing with moving average over 7 bins (350bp) was applied. Grey vertical lines indicate positions of ARS sequences in the region. Smoothing with moving average over 7 bins (350bp) was applied.

**Supplementary Figure 3.**
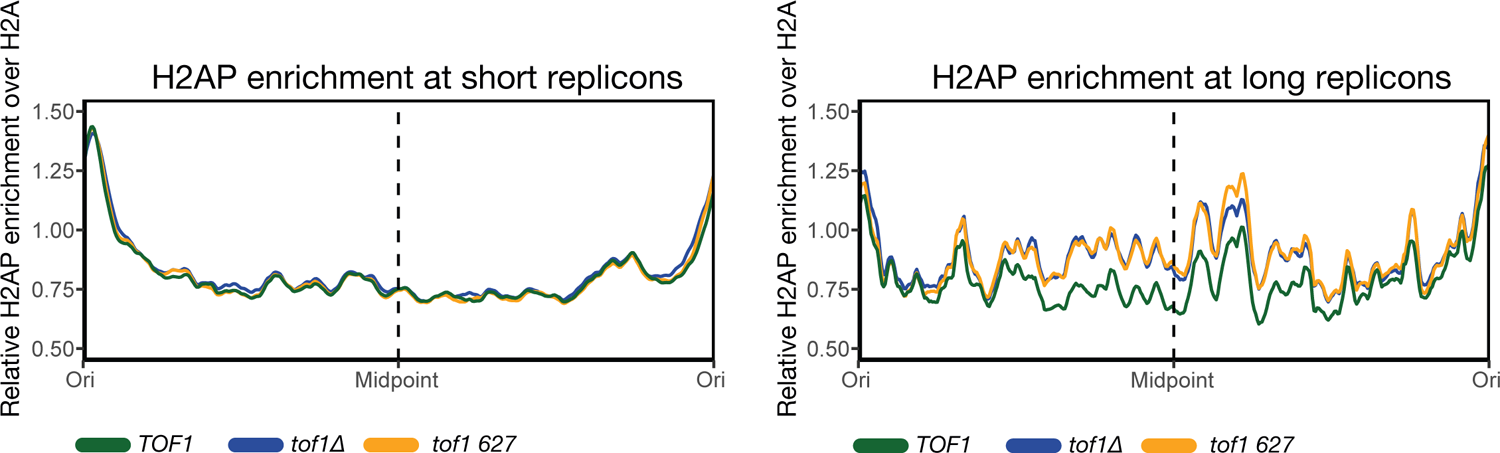
Cells expressing tof1 627 show similar accumulations of H2AP to TOF1 deleted cells across long replicons Relative H2AP enrichment in*TOF1wt*, *tof1Δ* and *tof1 627* cells over short replicons (20kb to 50kb) (left) and long replicons (>60kb) (right). Smoothing with moving average over 20 bins (1 kb) was applied.

**Supplementary Figure 4.**
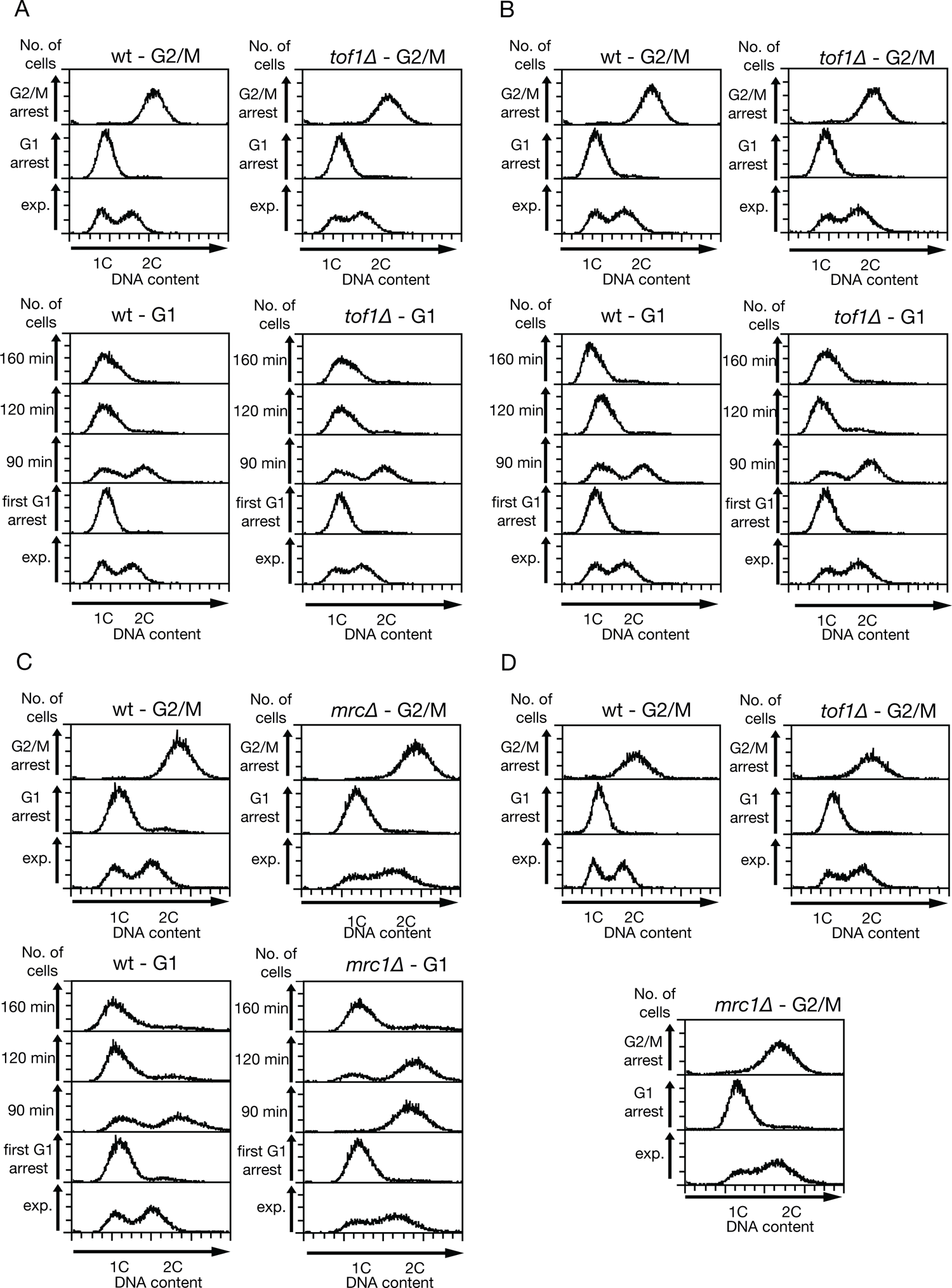
FACS analysis of cell cycle synchronization experiments shown in Figure 4 (A) G2/M (top) and G1 (bottom) arrest of *wt* and *tof1Δ* cells. Data used in Figure 5A. (B) repeat of (A) Used in Figure 5 AB. (C) G2/M (top) and G1 (bottom) arrest of *wt* and *mrc1Δ* cells. Used in Figure 5 AB. (D) G2/M arrest of *wt*, *tof1Δ* and *mrc1Δ* cells. Used in Figure 5AB.

**Table 1.**
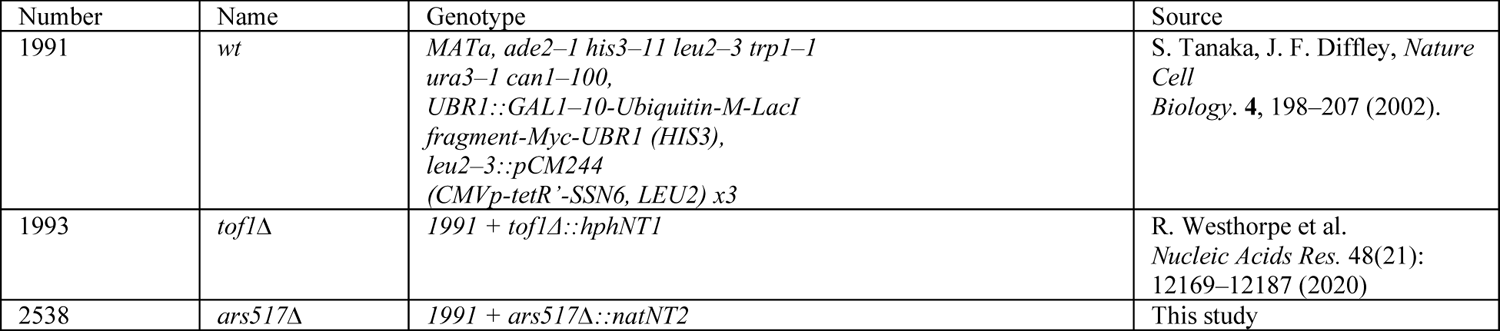
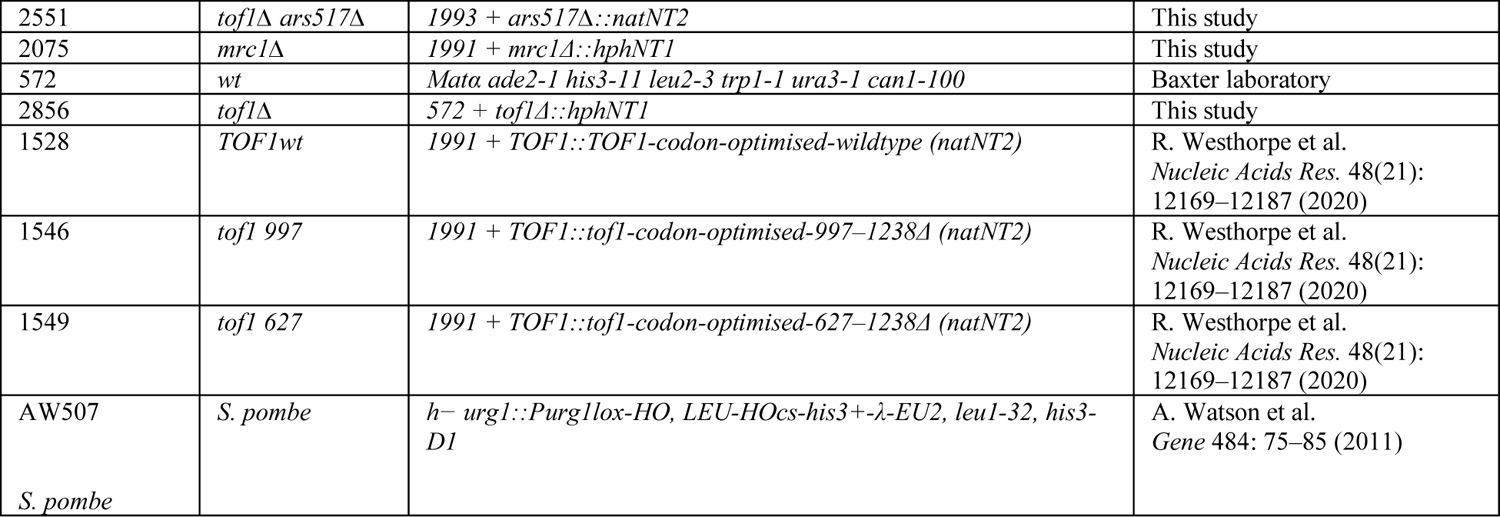
Yeast strains used in this study.

**Supplementary Table 1.**
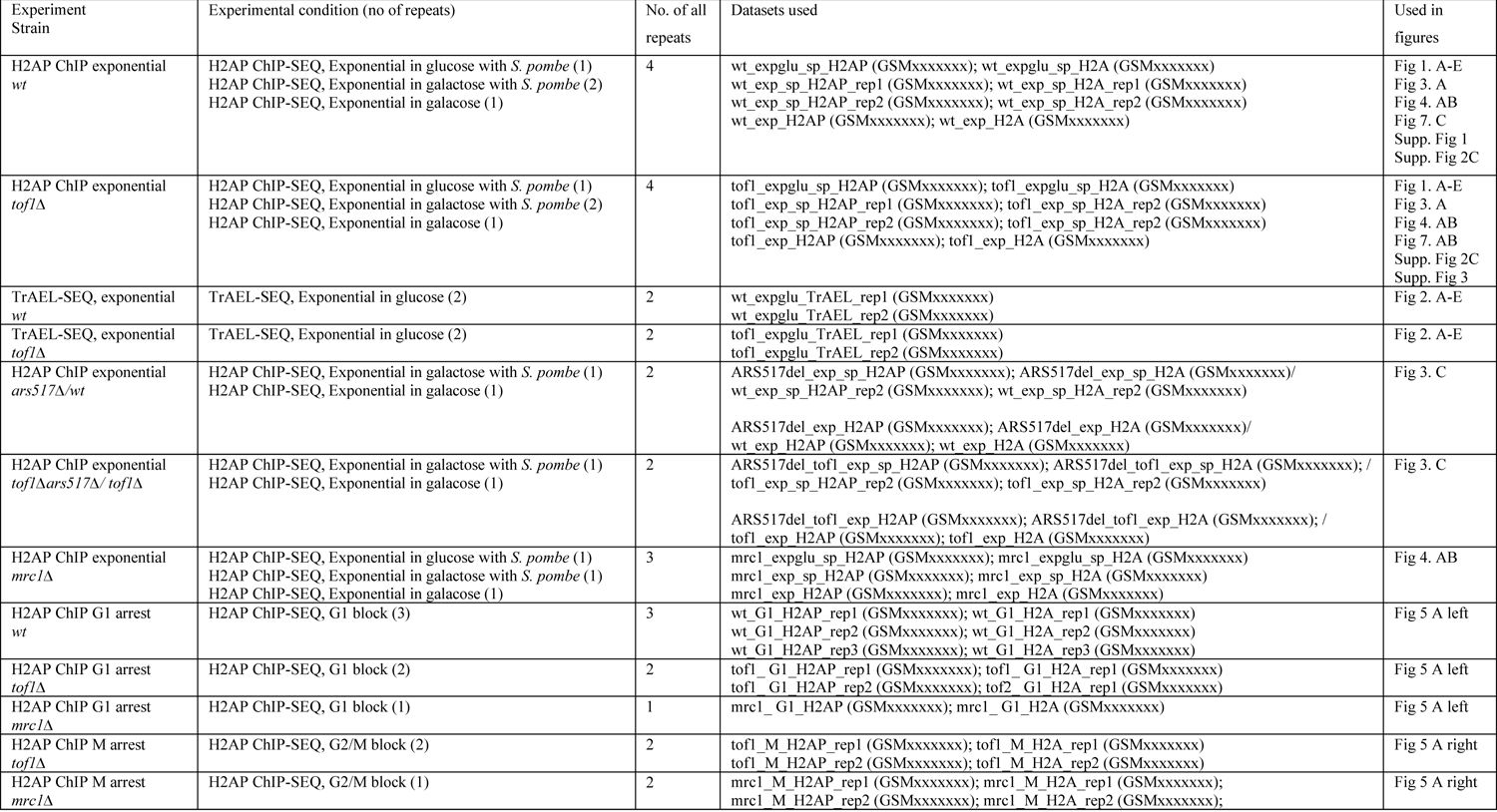
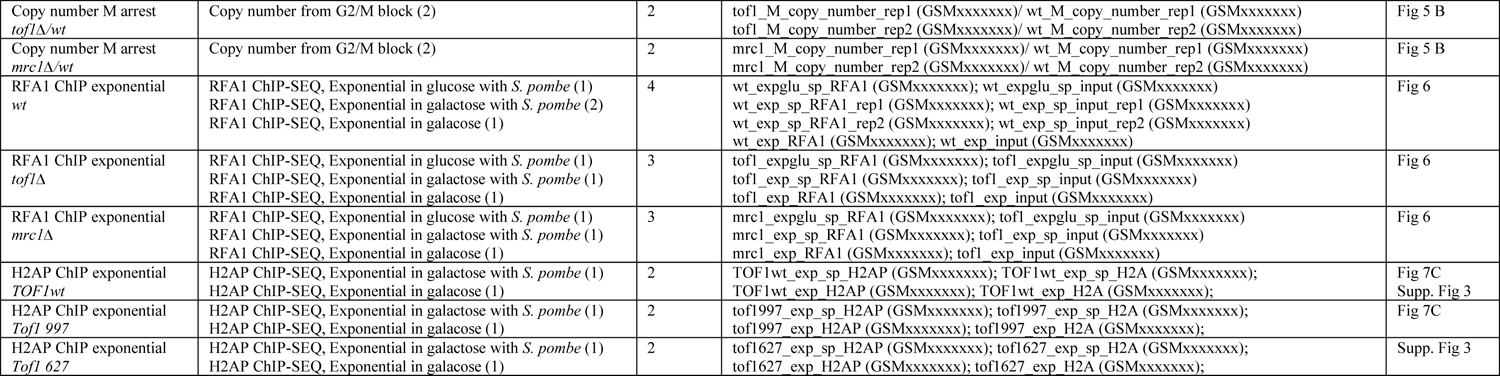
Number of repeats and experiment conditions used.

## Key resources table

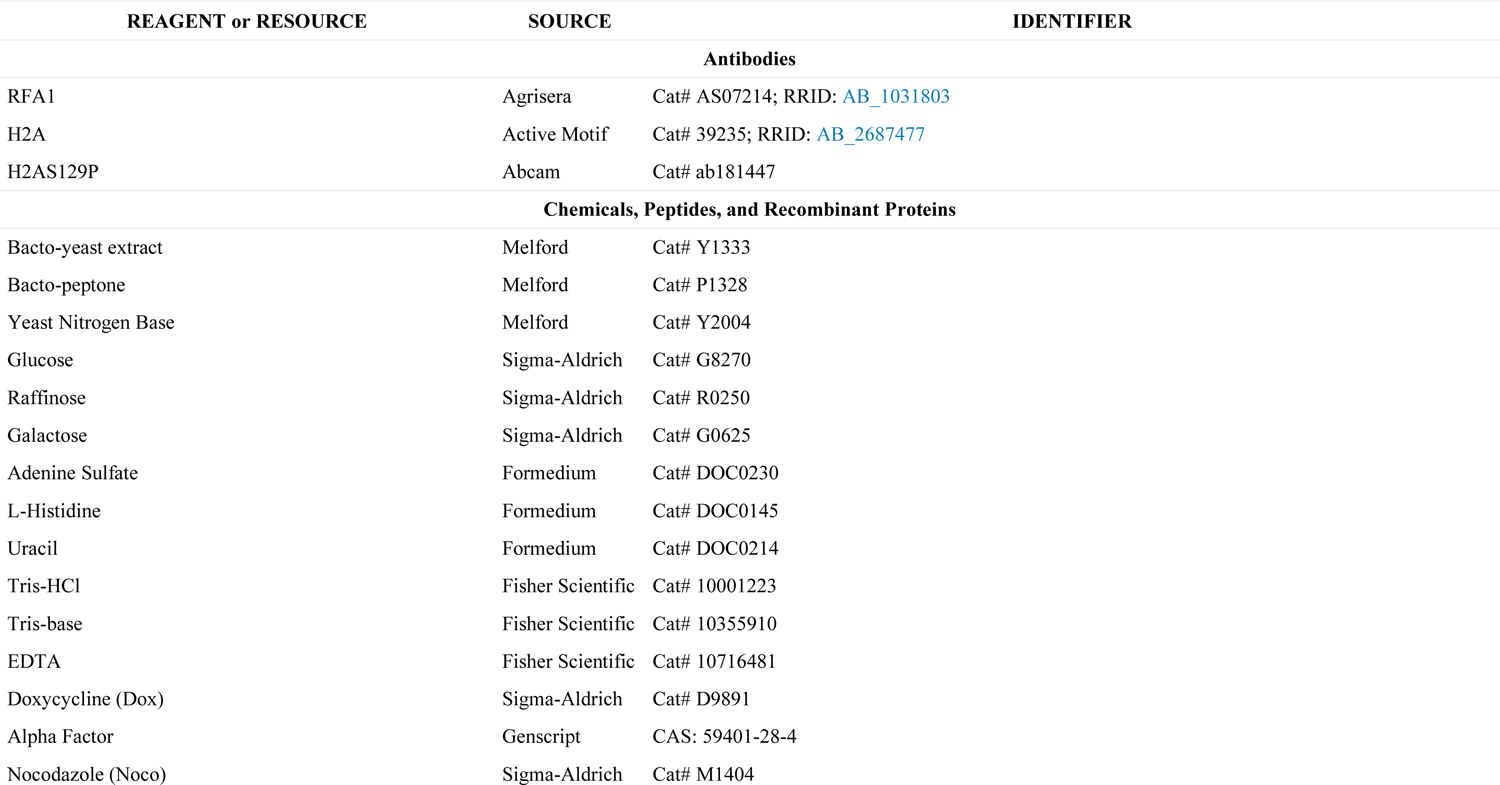
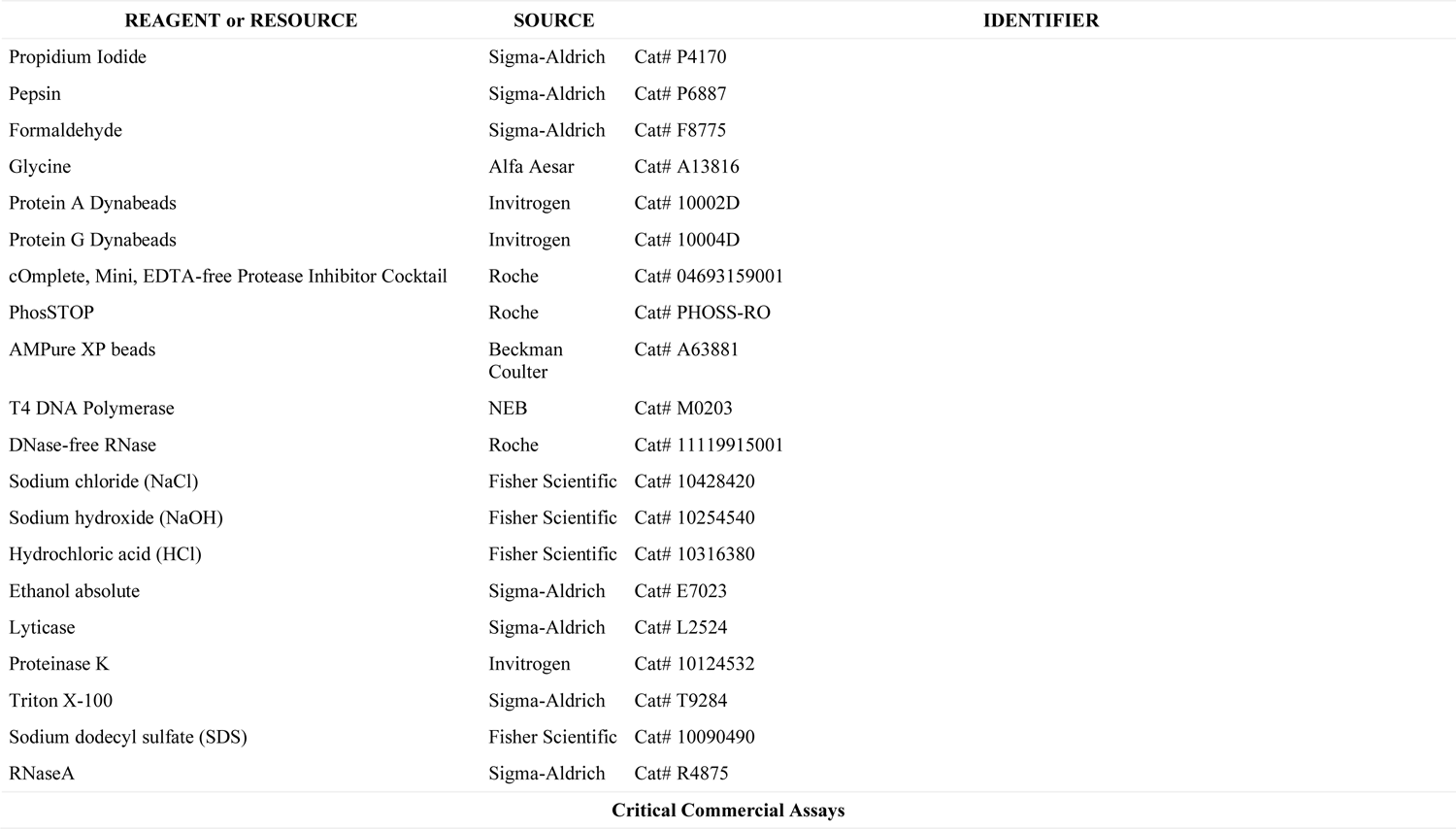
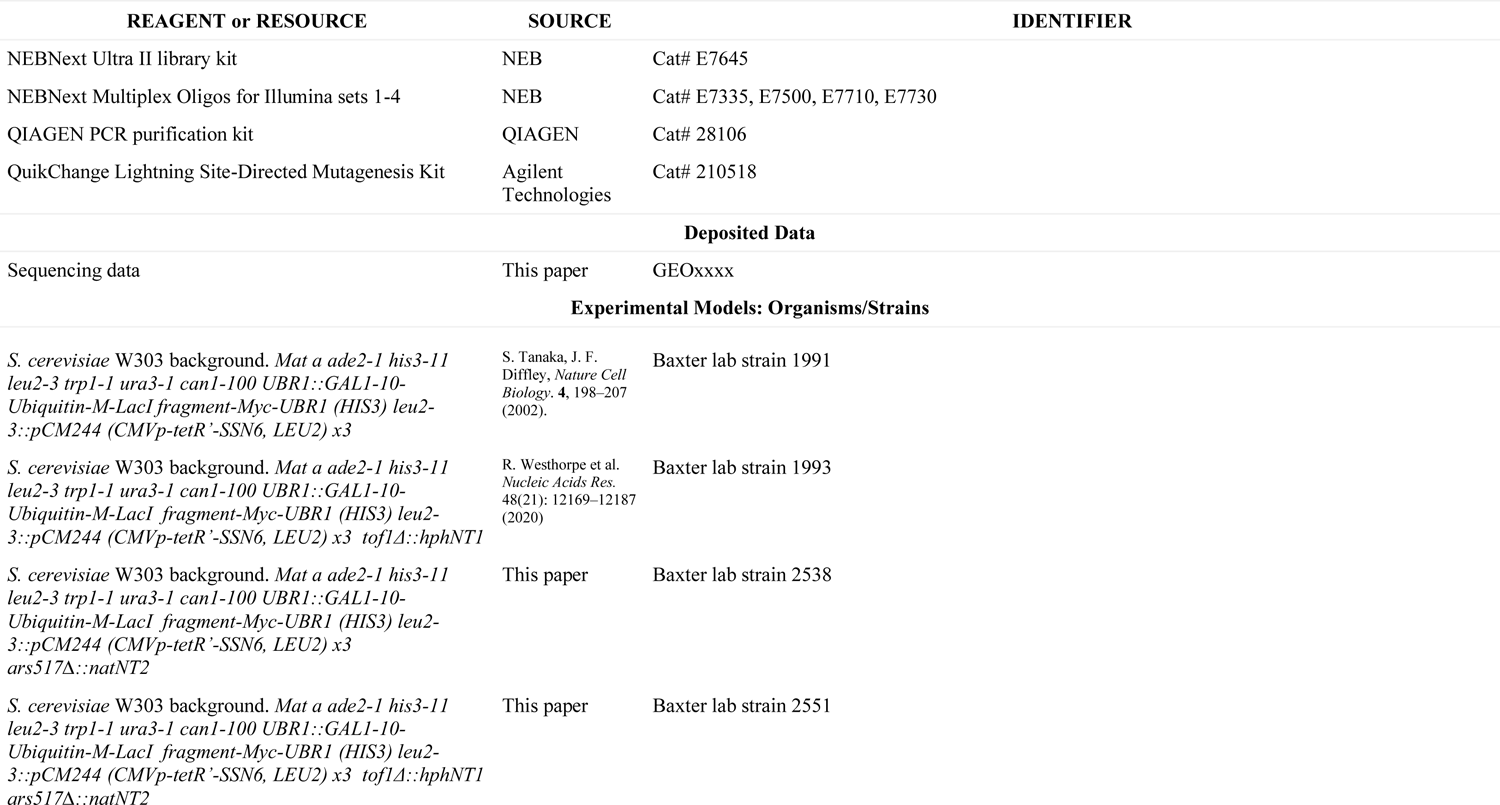
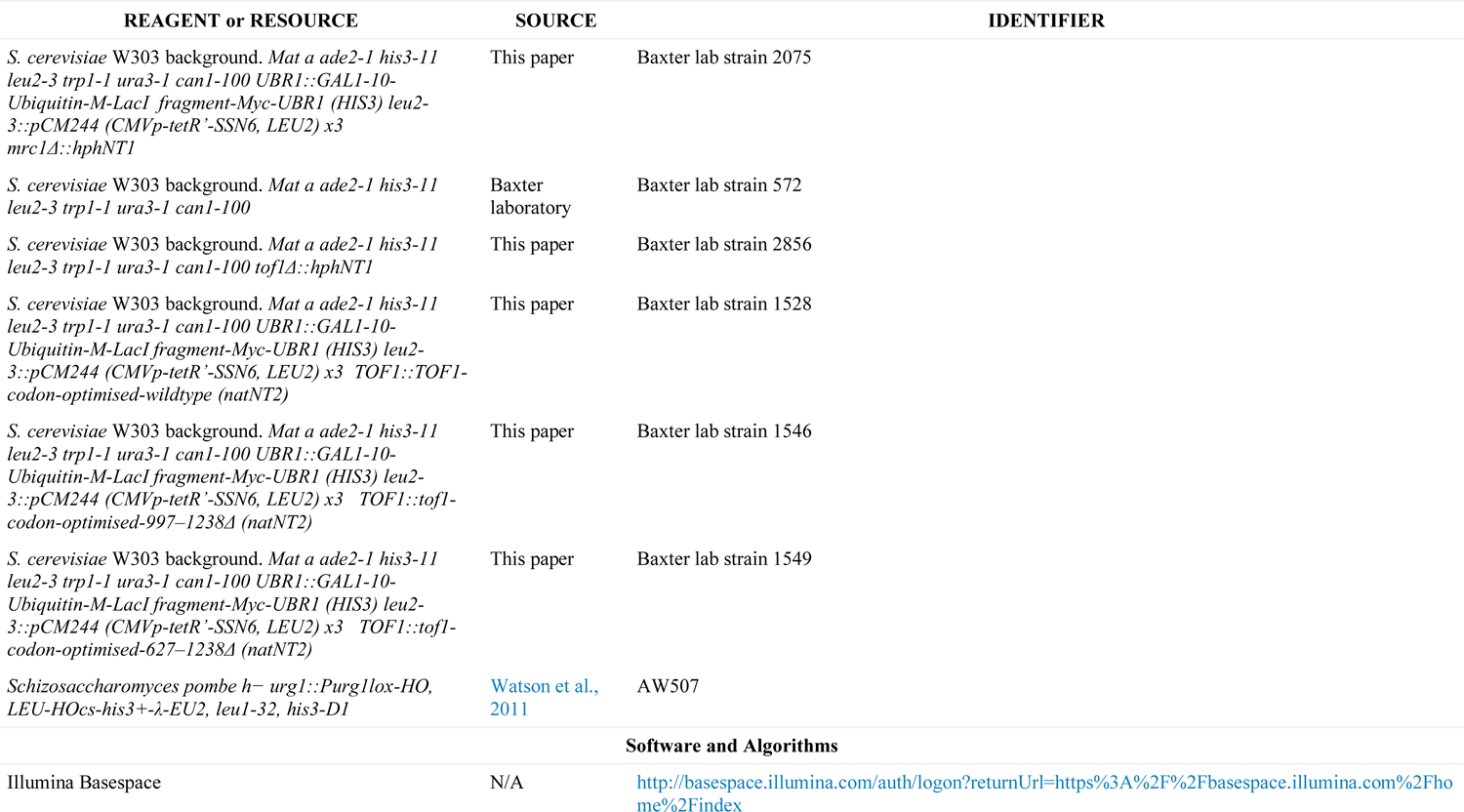
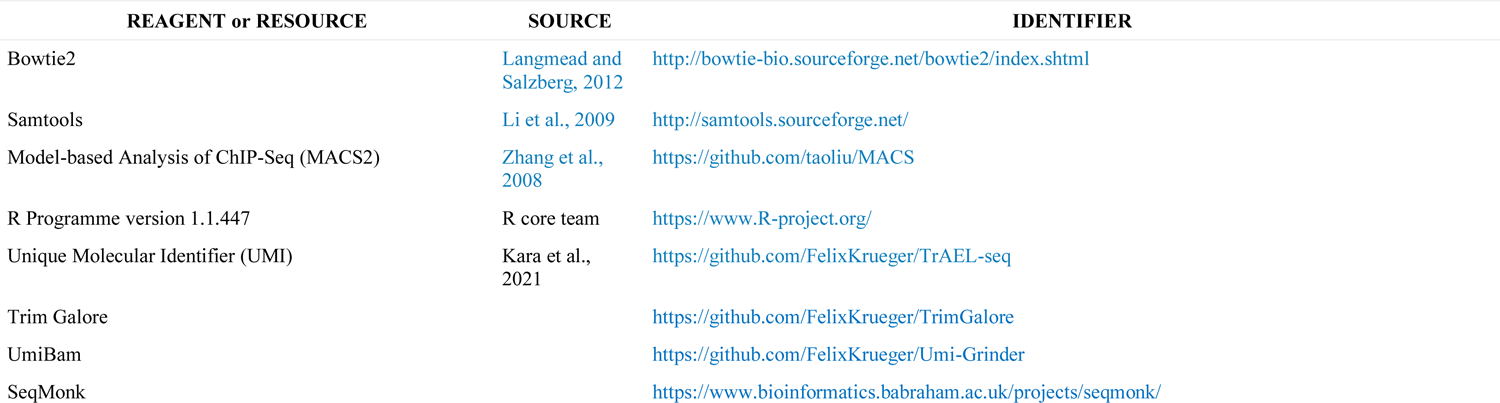
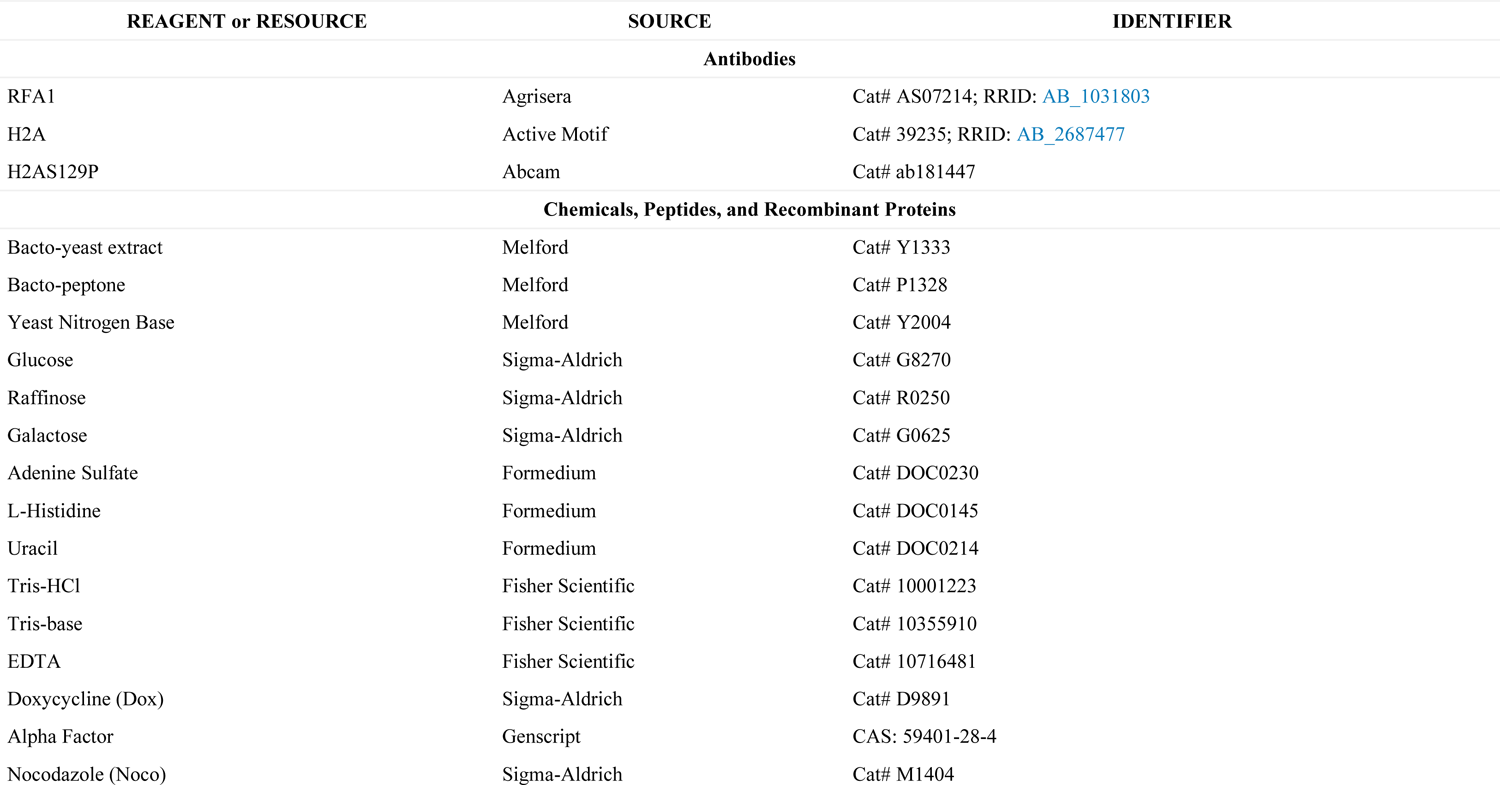
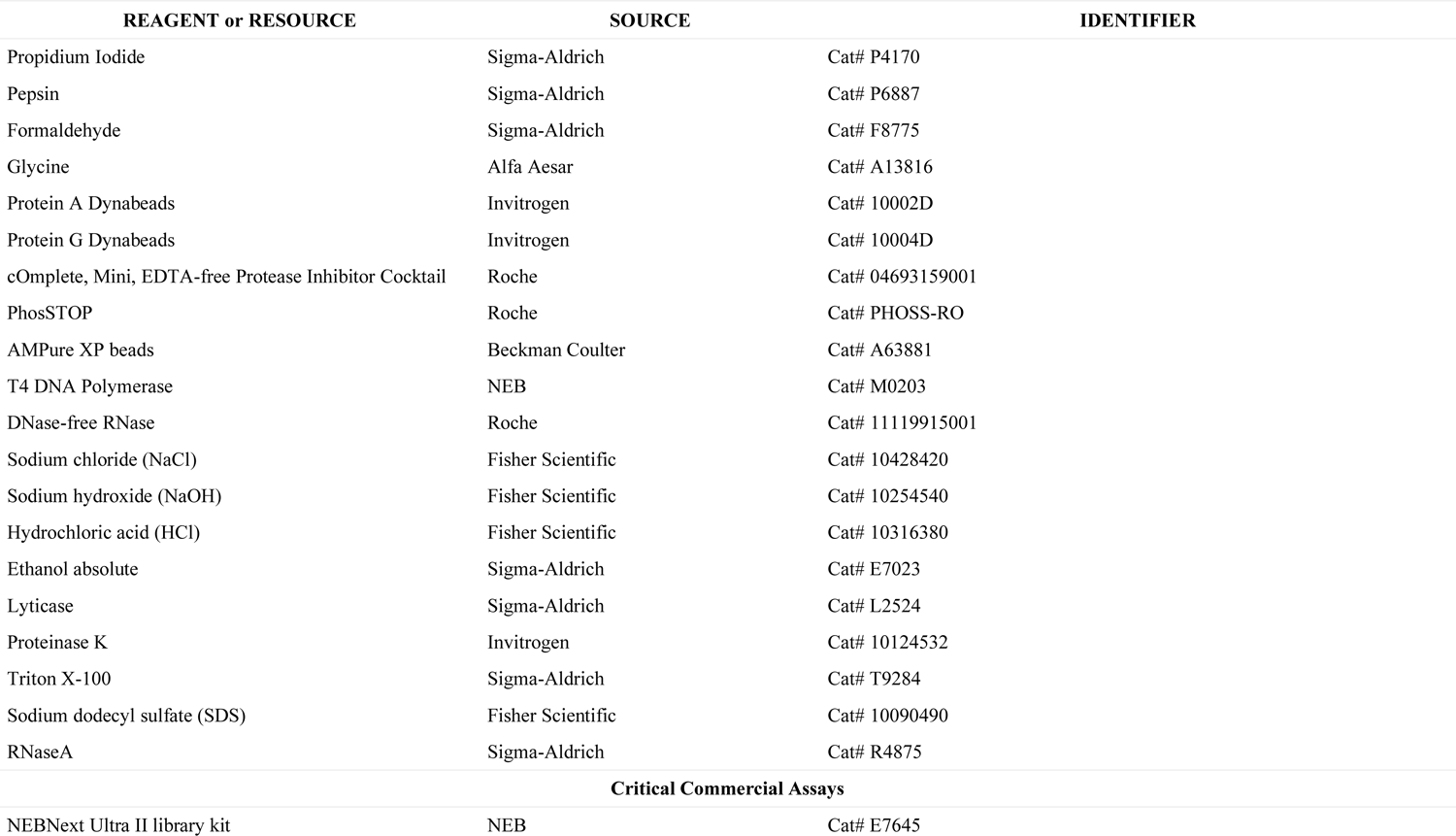
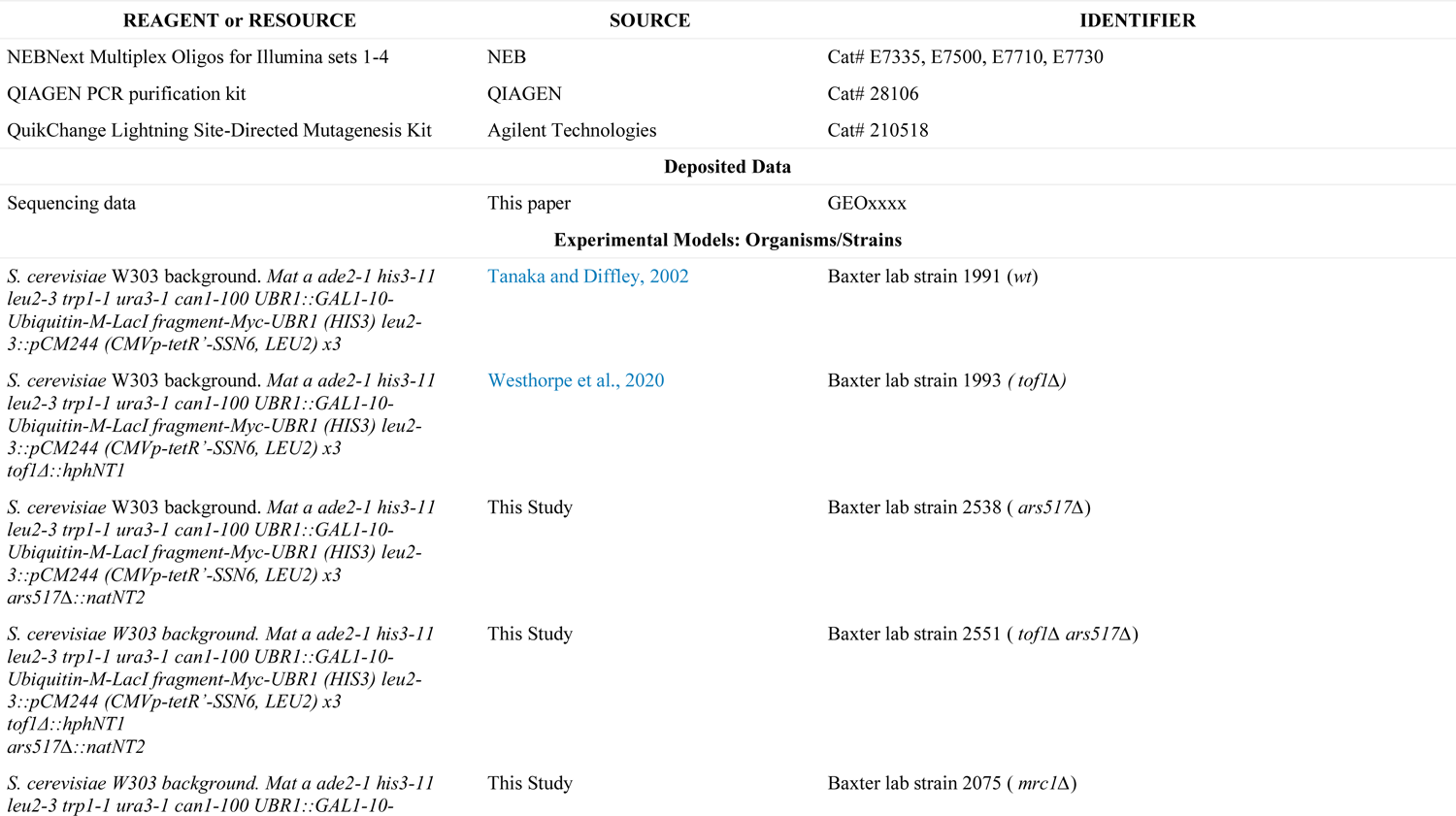
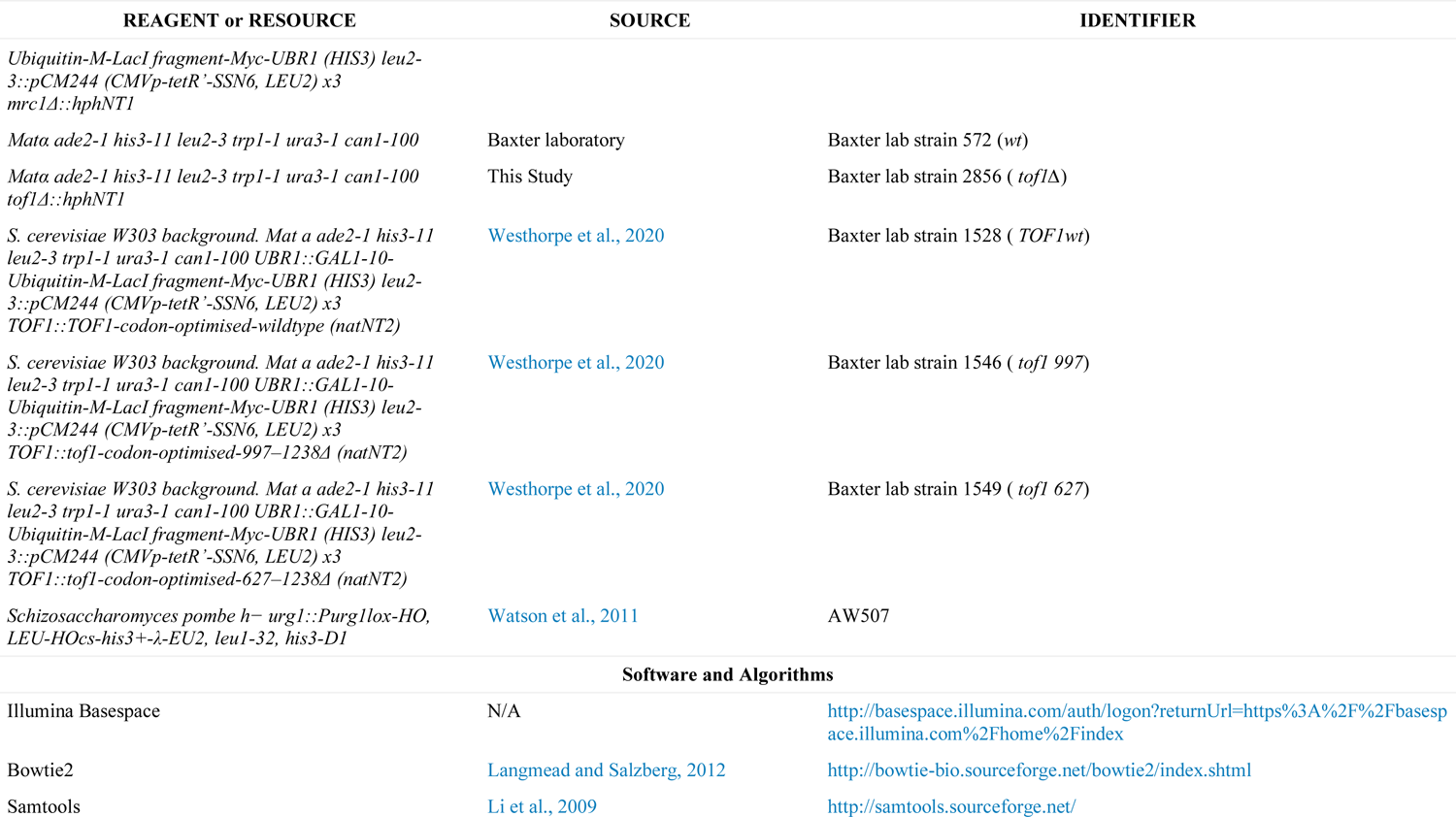
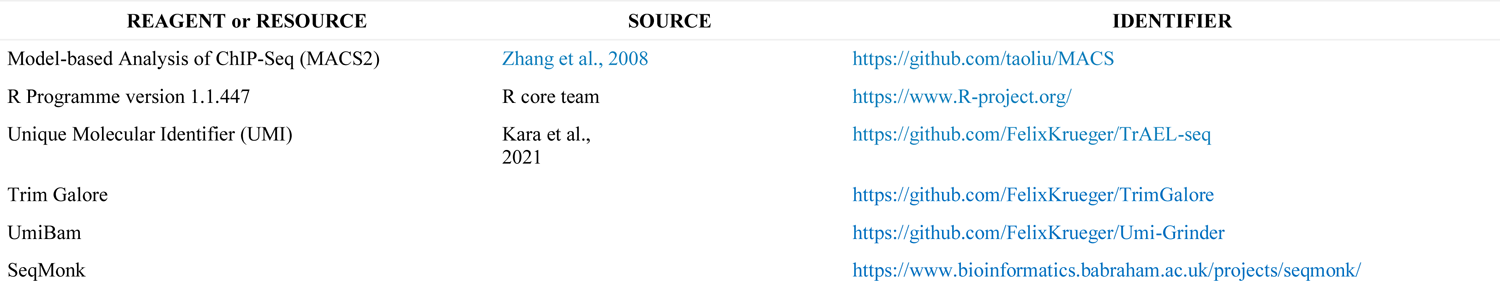

